# NMDA Receptors Control Activity Hierarchy in Neural Network: Loss of Control in Hierarchy Leads to Learning Impairments, Dissociation, and Psychosis

**DOI:** 10.1101/2023.01.06.523038

**Authors:** Yuxin Zhou, Jenn Lingshu Wang, Liyan Qiu, Jordan Torpey, Jemma Glenn Wixson, Mark Lyon, Xuanmao Chen

## Abstract

While it is known that associative memory is preferentially encoded by memory-eligible “primed” neurons, *in vivo* neural activity hierarchy has not been quantified and little is known about how such a hierarchy is established. Leveraging *in vivo* calcium imaging of hippocampal neurons on freely behaving mice, we developed the first method to quantify real-time neural activity hierarchy in the CA1 region. Neurons at the top of activity hierarchy are identified as primed neurons. In cilia knockout mice that exhibit severe learning deficits, the percentage of primed neurons is drastically reduced. We developed a simplified neural network model that incorporates simulations of linear and non-linear weighted components, modeling the synaptic ionic conductance of AMPA and NMDA receptors, respectively. We found that moderate non-linear to linear conductance ratios naturally leads a small fraction of neurons to be primed in the simulated neural network. Removal of the non-linear component eliminates the existing activity hierarchy and reinstate it to the network stochastically primes a new pool of neurons. Blockade of NMDA receptors by ketamine not only decreases general neuronal activity causing learning impairments, but also disrupts neural activity hierarchy. Additionally, ketamine-induced super-synchronized slow oscillation during anesthesia can be simulated if the non-linear NMDAR component is removed to flatten activity hierarchy. Together, this study develops a unique method to measure neural activity hierarchy and identifies NMDA receptors as a key factor that controls the hierarchy. It presents the first evidence suggesting that hierarchy disruption by NMDAR blockade causes dissociation and psychosis.

## Introduction

It is known that a small portion of sparsely distributed neurons with elevated excitability in memory-related regions are preferentially allocated to encode associative memory information ^1–5^. A long-held engram theory posits that memory-eligible neurons “activate” during learning to acquire a memory and the same group of neurons “re-activate” during recall to retrieve the memory ^3,4,6,7^. While the “activation” and “re-activation” model has greatly advanced our understanding of contextual fear memory and spatial memory ^7–12^, it has limitations in explaining the network basis of associative memory that does not critically depend on spatial or contextual information. First, by detecting the expression of immediate early genes such as c-Fos ^3^, the “neuronal activation” captures the cellular step of memory consolidation or the “nuclei activation” of memory-eligible neurons ^3^, rather than the exact process of memory acquisition. Second, while neurons with high excitability are preferentially recruited to acquire or retrieve a memory ^13–17^, the same group of neurons can maintain a high-activity status for several days ^18–20^. If simply the activation of these neurons encoded a memory, it could not allow for the same subset of neurons to encode different memories within the same period. Third, our recent *in vivo* calcium imaging combined with trace fear conditioning study has demonstrated that primed hippocampal neurons, which are already randomly active prior to training, develop burst synchronization when an associative trace memory is being formed or retrieved ^20^. We further found that it is not the “activation”, but rather the burst synchronization of primed neurons that is crucial for associative learning ^20^. Consistently, repetitive aversive stimuli hardly activate relatively “silent” hippocampal neurons to actively engage in associative memory formation ^20^. Because there is a neural activity hierarchy in memory-related regions ^14,21,22^, the flow of neural information is preferentially transmitted from primed neurons to primed neurons in the network. We reason that maintaining a dynamic neural activity hierarchy is vital for associative learning and sustaining mental health. Therefore, elucidating the mechanisms of how neural activity hierarchy is established and maintained will not only advance our understanding of associative memory formation, but also unveil the pathophysiology of multiple cognitive dysfunction-related mental disorders, including schizophrenia.

The N-methyl-D-aspartate (NMDA) receptors (NMDARs) represent one of the most important drug targets and ionotropic receptors that mediate the excitatory neurotransmission in the nervous system ^23^. The receptors are best known to mediate multiple forms of synaptic plasticity including long-term potentiation ^24–26^, a cellular mechanism widely thought to be essential for learning and memory formation ^24,27–29^. Malfunction of NMDA receptors is associated with a broad spectrum of brain disorders, varying from Alzheimer’s disease ^30^ and schizophrenia ^31^, to neuropathic pain and excitotoxicity ^32^. Research on NMDA receptors and the receptors-mediated signaling pathways has been very active for over 40 years^33–36^, to determine the molecular mechanisms of neurodevelopment, synaptic plasticity, and excitotoxicity, and to explore whether pharmacological modulation of NMDA receptors treats dementia, clinical depression, and other brain disorders. To date, numerous agents have been identified to target NMDA receptors, including ketamine, nitrous oxide, dextromethorphan, memantine, methoxetamine, and MK-801 ^23,37,38^, of which, some are used clinically to treat Alzheimer’s disease, pain, or excitotoxicity-related disorders ^38–40^. The best-known agent is ketamine, which is used as an anesthetic and fast-acting antidepressant ^41,42^. However, high-dose drug usage or high-potency blockade of NMDA receptors often leads to dissociation, hallucination, cognitive impairments, or psychotropic effects in patients and individuals who abuse these agents ^41,42^. For this reason, ketamine is often utilized as a pharmacological tool in schizophrenia research ^31^. Consistently, hypofunction of NMDA receptors is associated with schizophrenia in mouse models and humans ^31,43,44^. However, it remains to be elucidated how NMDA receptor blockade by ketamine or other agents causes dissociative and psychotic symptoms, and why hypofunction of NMDA receptors is closely linked to schizophrenia.

While certain proteins such as CREB ^15,45^ and potassium channels ^22,46,47^ are found to regulate intrinsic neuronal excitability affecting learning, memory formation or retrieval, a real time neural activity hierarchy has not been quantitatively determined previously. Leveraging deep-brain *in vivo* calcium imaging acquired during freely behaving trace fear conditioning experiments, we sought to develop a method to measure neural activity hierarchy and quantitatively establish primed neurons that engage in trace memory formation. We have identified two criteria that are the most relevant to distinguish primed neurons from non-primed neurons: (1) high basal activity levels prior to training with increased activity through training; and (2) the ability to form burst synchronization during trace fear conditioning and recall testing. We included these criteria in our analysis to develop a novel numerical method that quantifies *in vivo* neural activity hierarchy.

To understand the mechanisms of how a neural activity hierarchy is established in neural network, we developed a simplified neural network simulation model that consists of a two-dimensional array of neurons. It incorporates simulation of synaptic transmission that has linear and non-linear weighted synaptic components, which simulate the AMPAR- and NMDAR-mediated conductance, respectively. One crucial difference between the AMPAR- and NMDAR-mediated conductance is the voltage-dependent magnesium-blockade of the NMDAR^48,49^. In addition to glutamate and glycine binding, NMDAR requires the depolarization of membrane potential to relieve the magnesium blockade to fully open the channel for ion flow ^49,50^. This means that NMDARs in active (depolarized) neurons are more responsive to glutamate stimulation than in inactive neurons (less depolarized), thereby generating a non-linear ionic conductance for the excitatory synaptic transmission^49,50^. This scenario favors efficient synaptic communication to active neurons over inactive neurons. Consequently, NMDARs in active neurons contribute to a higher portion of postsynaptic currents than in inactive neurons, even if excluding the secondary effect of synaptic plasticity. Our simulation reveals that the presence of the non-linear conductance is required for establishing neural activity hierarchy and a moderate ratio of non-linear to linear conductance naturally leads to the emergence of primed neurons. We further found that the basal neuronal activity levels modulate the development of neural activity hierarchy and a high initial activity level speeds up the development of neural hierarchy. To experimentally evaluate the role of NMDARs in controlling neural activity hierarchy, we used ketamine to block NMDAR during trace fear conditioning, in conjunction with *in vivo* calcium imaging study. We discovered that NMDAR blockade not only decreases overall neuronal activity leading to learning impairments, but also disrupts neural activity hierarchy. In addition, ketamine, as a dissociative anesthetic, induces EEG waveform that manifests high-amplitude slow oscillation in the neocortex and hippocampus ^51,52^. Interestingly, this waveform can be nicely modeled by our neural network simulation if the non-linear conductance is removed. Together, we show for the first time that the non-linear synaptic conductance mediated by NMDARs is crucial for establishing and maintaining neural activity hierarchy and NMDAR blockade disrupts neural activity hierarchy, leading to learning impairments, dissociative, or psychosis-like behaviors.

## Methods

### Mice

All animal-related procedures were approved and conducted in accordance with the guidelines of the Institutional Animal Care and Use Committee of the University of New Hampshire. Mice were maintained on a 12-h light/dark cycle at 22°C and had access to food and water ad libitum. The Ift88 floxed mouse strain ^53^ was cross-bred with UBC-Cre/ERT2 as previously reported (C57B1/6j genetic background, mixed sexes) ^54^. At the age of 8-10 weeks, Ift88 flox/flox: UBC-Cre/ERT2 mice were orally administrated with tamoxifen (0.2 mg/g body weight, consecutive 7 days) to induce Cre recombinase expression to ablate Ift88 and generate Ift88 cilia KO mice, or with corn oil to make vehicle control mice ^55^. Ift88 flox/+ (or Ift88 +/+) UBC-Cre/ERT2 mice with the same tamoxifen treatment were used as genotype controls. We observed that vehicle control and genotype control mice exhibited very similar results, and their data were combined into one control group.

### Mice surgery and AAV viral vector injection

Surgery was performed one week after tamoxifen/vehicle administration, as previously described ^20^. Mice were anesthetized by 1.5-2% isoflurane and mounted on a stereotaxic frame (David Kopf Model 940). A special canula implant (Mauna Kea Technologies, Paris, France) was installed above a small window through the skull (coordinate: AP: −1.95 to −2.05 mm relative to the Bregma, ML: −1.6 mm relative to the midline). The cannula implant allowed the injection of viral vector (0.5 µl AAV1-Syn-GCaMP6m, Addgene, ID 100841) into the hippocampal CA1 region (coordinate: DV: −1.45 to -1.55 mm from the skull) to express calcium indicator and allow a fiber-optic imaging microprobe to go through. Mice were used for *in vivo* calcium imaging and trace fear conditioning 7-8 weeks after the canula implantation surgery.

### Imaging acquisition

*In vivo* calcium dynamics were acquired by a Cellvizio Dual-Band 488/660 imaging system (Mauna Kea Technologies, Paris, France) using laser excitation wavelength at 488 nm (maximum power output < 22 mW), fluorescence detection wavelength between 502-633 nm. A CerboflexJ NeuroPak deep-brain fiber-optic microprobe (having over 7600 optical fibers, lateral resolution 3.3 μm, maximum field of view ∅ = 325 µm) collected the green florescent signal of GCaMP6m from hippocampal CA1 region (coordinate relative to the Bregma: AP: −1.95 to −2.05 mm, ML: −1.6 mm, DV: −1.75 to -2.55 mm). GCaMP6m fluorescent signals were acquired every training or recall cycle to monitor the real-time calcium dynamics. Imaging sessions were also recorded prior to training and recall, after training, during resting and under isoflurane-induced anesthesia. Acquired imaging data (25 Hz, 66 seconds for training cycles, 55 seconds for others) were analyzed off-line using IC Viewer 3.8 (Mauna Kea Technologies, Paris, France). Regions of interest (ROI) of the calcium signals for individual cells were manually selected to cover at least 3 fibers. The ROIs with intensity lower than 400 Analog-Digital Unit (ADU) in any recorded cycles were excluded from analysis. ROIs from background areas covering 50 fibers without GCaMP6 expression were selected to detect the background pattern, which, in turn, was used to eliminate imaging noise and artifacts. 70 -140 individual cell ROIs and 12 background ROIs were collected from every animal. ROIs also contain X-Y coordinate information, which allows for plotting neurons’ locations in imaging fields. Representative raw calcium traces throughout the manuscript are the calcium intensity relative to the basal level (%ΔF/F) computed as the GCaMP6m fluorescence intensity divided by the intensity average of 1 second (25 frames) before tone for every recording. To present coherent bursting among primed neurons, we filtered imaging data using a 1 Hz high-pass Fourier filter. The high-pass filter was implemented by first subtracting a low-degree polynomial projection of the data and then using a Fourier transform and directly zeroing frequency lower than 1 Hz. To compute synchronization level and perform other analysis, imaging data were first transformed to 5-second moving SD traces and the background artifacts were removed.

### Trace fear conditioning test

Trace fear conditioning was carried out as previously described ^20,55^, but in conjunction with *in vivo* calcium imaging. Behavioral tests was performed 6-8 weeks after AAV-GCaMP injection and canula implantation. Mice carrying imaging probes were placed in a fear conditioning chamber (Med Associates Inc, Vermont). Following 10 minutes of exploration and acclimation, a neutral tone (3 kHz, 80 dB, 15 s) served as the conditioned stimulus (CS), followed by a mild electric foot shock (0.7 mA, 1 s) as the unconditioned stimulus (US). Tone and foot shock were separated by a 30-second trace period to prevent direct CS-US association. The training cycle was repeated 7 times to develop a trace fear memory. Behavioral data were recorded for another 20 minutes after the 7th cycle to monitor post-training behaviors. After training, mice were placed in a resting box with ad libitum bedding and food. After 2-3 rest, memory retrieval testing was assessed in a novel environment (solid plastic walls and floor) following 10 minutes of acclimation. A tone (3 kHz, 80 dB, 5 s) served as the conditioned cue, with recall cycles (10 s pre-tone, 5 s tone, 30 s delay, no shock, 10 s post-tone, and 110 s interval) repeated 5 times. Training and recall sessions were videotaped (25 frames/second), and mouse behaviors were analyzed using Noldus EthoVision XT to measure locomotion, freezing, and velocity. Freezing behavior was defined as real-time velocity below 1 mm/s, as automatically determined by EthoVision XT.

### Background noise elimination

The lightweight of the fiber-optic imaging microprobe permits animals to behave freely. However, free behaving imaging may generate certain motion artifacts and background noise. To detect background imaging noise, 12 background ROIs were selected from areas without GCaMP6 expression. Any consistent fluctuations in these background ROIs were considered imaging noise or motion artifacts. To remove any background noise patterns, the calcium traces from the background regions were first transformed by computing a standard deviation over a 5-second moving window. While the SD trace over a 5-second window does “smear” the data over those 5-seconds, this has the benefit of reducing the effect of any white noise in the data in addition to eliminating potentially spurious phase information. The final calculations are not particularly sensitive to the precise choice in the length of the window as long as the same process and window lengths are used for both the background regions and the individual neurons. The background 5-second SD traces were normalized and then analyzed through a principal component analysis (PCA) analysis, which is used to identify any patterns consistent among the background regions. The principal components in the PCA analysis with scores that were above the average (approximately 4-5 patterns) were then deemed to be artifacts in the data and were used as the background artifact/noise patterns. The identified artifacts were then eliminated from all individually neuronal calcium signals’ 5-second SD traces by calculating the projection of the individual signal patterns onto the identified artifact patterns and subtracting that projection from the individual neuron’s SD trace, thus orthogonalizing all the individual neuron’s SD traces to all artifact patterns.

### PCA analysis and the calculation of neuronal synchronization levels

Once the 5s-moving SD trace of each individually measured neuron had been computed and all artifact patterns removed, a second PCA was performed to determine the major pattern in the individual neuron responses. Here we looked at the singular vector with the largest singular value (or the principal component with the largest score) which represents a signal pattern that was exhibited more often in the neurons than any other pattern. Since this major pattern was constructed after the SD traces of all the individual neurons have had any background artifacts removed, the major pattern is orthogonal (or completely uncorrelated) to any of the identified background artifacts patterns.

After the major pattern for neuronal signals has been identified, correlations between the SD traces of all individual neurons and the major pattern can be computed. Similarly, the correlation between any given pair of SD traces can be computed in the heatmaps. Values for these correlations which are above 0.7 signify strong correlations ^56^. When the principal component score corresponding to the major pattern was significantly larger than the next largest score, many individual SD trace signals were found to have high correlations in our imaging data. These high correlations with a single major pattern are a clear indication of neuronal priming. The extent to which the neuron is primed can be estimated by the number of neurons exhibiting a strong correlation with the major pattern.

We collected calcium images of individual neurons under different conditions, including under anesthesia using 2% isoflurane. The neuronal calcium dynamics under anesthesia served as the internal reference for normalization to eliminate individual variance in signal detection and GCaMP expression and to allow for quantitative comparison between different animals. The activity levels of individual neurons were calculated as the ratio of the SD-of-SD value during training cycles to the SD-of-SD value under anesthesia. In our previous report ^20^, the neuronal activity levels were estimated using the ratio of variances of the calcium traces during training relative to those under sleep. That method could distinguish silent neurons from primed neurons clearly but was unable to accurately separate intermediately active neurons from primed neurons. We define primed neurons as high-activity neurons with strong responses to learning cues and able to form burst synchronization with each other. Some intermediately active neurons can exhibit relatively high activity but while their calcium traces showed consistent high amplitude, they were not as responsive to learning cues in that their SD traces remained relatively uncorrelated with the major pattern. The second order activity level measured by the SD-of-SD more accurately separates primed neurons from intermediately active neurons than the first order activity level measured by variance. Using SD-of-SD highlighting bursting, we can differentiate bursting signals from consistent high-amplitude fluctuations and efficiently distinguish primed neurons from intermediate and silent neurons.

### Immunofluorescence staining

Mice were euthanized, and brain tissues were extracted and fixed with 4% paraformaldehyde at 4°C overnight. After sliced, brain samples were subjected to fluorescence immunostaining using primary antibodies against GFAP (1:500, Cat# Z0334, Dako), and c-Fos (1:500, Cat# MCA-2H2, EnCor), followed by secondary antibodies (conjugated with Alexa Fluor 488, 546, or 647, 1:500) to verify GCaMP6 expression, imaging site and c-Fos expression, reactive astrogliosis (GFAP staining) in the hippocampal CA1 region. The number of c-Fos positive neurons normalized to the nuclear number and GFAP/DAPI intensity ratio were compared between the surgical and non-surgical hemispheres in 12 slices collected from 4 animals. All fluorescence images were captured using a Nikon A1R HD confocal microscope acquired using the same settings including exposure time, gain and laser power settings. Images were processed and analyzed by Fiji ImageJ ^57^.

### Bursting analysis of time-frequency plots of imaging

While SD traces only capture the magnitudes of fluctuations in the signal, some information on the frequency of the fluctuations can be obtained. The maximum frequency that can be directly observed is limited to 12.5 Hz (due to the 25 Hz sampling rate) while any higher frequencies will be aliased and appear as a frequency from 0 to 12.5 Hz. Time-frequency plots were generated by computing short-time Fourier transforms using the “stft” algorithm in MATLAB. The fast Fourier transform (FFT) length was set to 512 with an overlap of 252 and a 256 length Kaiser-Bessel-derived window with a parameter beta of 10. The frequency as a function of time can then be displayed as a heat map. Synchronization and bursting patterns can still be observed even if the frequency data is aliased and the precise frequencies in the signal exhibiting the bursting behavior cannot be determined.

### Neural network simulation

A two-dimensional computational array of *N*_r_ = 2,000 rows (cell number per layer) by *N_c_* = 30 columns (or layers) of neurons was simulated through 12,500 computational iterations. The model serves as a relatively simple demonstration of a possible mechanism for neuronal priming. As the precise configuration of a grouping of neurons in the brain is extremely complex, many simplifying assumptions were made to make the simulation tractable, though extensions and generalizations of this model will be explored in future work. The parameters used in the model were modified through trial and error to obtain the results shown and using different parameters can yield significantly different results and distributions, although alternate parameters can also yield similar results.

Several properties were kept track of for each neuron at each time cycle. The activation level of the neuron in the *i*^th^ row and the *j*^th^ column is labeled *A_ij_*, representing the activation level of the neuron. The net charge in the neuron was scaled to a unit interval so that if *A_ij_* reaches or exceeds the value of 1, the neuron will fire, which will result in neurotransmitter release and signals being received by neurons in column *j* + 1. The value *F_ij_* is the amount a given neuron has fired over approximately the last 2,500 time cycles and represents the recent firing rate of this neuron. To accelerate the simulation, *F_ij_* was updated only once every 50 iterations. This simulation was initialized with all values of *A_ij_* randomly and uniformly selected from the interval [0,1] and all values of *F_ij_* set to zero. A given neuron in layer *j* is assumed to be connected to all the neurons in layer *j* + 1 (that is when a neuron in layer *j* fires, the activation level of all 2000 neurons in layer *j* + 1 will increase to varying degrees). The connection strength, *S_ki_* between a neuron in layer *j* and row *i* to a neuron in layer *j* + 1 and row *k* is independent of *j* and given by

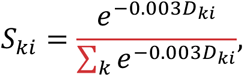

Where *D_ki_* is given by the minimum of three quantities |*k* − *i*|, |*k* − *i* − *N_r_*|, and |*k* − *i* + *N_r_*|. These definitions enforce a periodic boundary condition so that the bottom row is considered to be just above the top row. While a given neuron in layer *j* does connect to all neurons in layer *j* + 1, *S_ki_* decays rapidly as the rows *k* and *i* become further separated and so each neuron in layer *j* will only have a strong connection with a relatively small number of neurons in layer *j* + 1 (less than 200).

At the beginning of each cycle, 120 neurons (6% of 2,000) were randomly selected from the first layer and the activation level (*A_i_*_1_) of those 120 neurons were set to 1. Then, layer by layer, the entire mesh is checked to see if *A_ij_* ≥ 1, and all the neurons layer *j* with an activation level greater than or equal to 1 are fired and release signals to all connected neurons in layer *j* + 1, before moving on to check-up of the next layer *j* + 1.

As a neuron fires in layer *j* and row *i*, the transmission of signal to layer *j* + 1 is assumed to consist of three major components: *L_ij_* representing AMPAR-mediated synaptic conductance which is assumed to be linearly dependent on the on the firing of pre-synaptic neuron, *N_ijk_* representing NMDAR-mediated synaptic conductance taken to be non-linearly dependent on the firing of the post-synaptic neuron in layer *j* + 1 and row *k*, and *G_ij_* which mimics GABA_A_R inhibition. The inhibition is taken to be correlated to the firing activity of nearby interneurons which in turn are assumed to be correlated with the firing activity of the pre-synaptic neuron. Thus, *G_ij_*, just as *L_ij_*, is taken to be linearly dependent on the firing of the pre-synaptic neuron but of opposite effect.

When the neuron in row *i* and layer *j* fires, the activation levels for the neurons, *A_k_*_(*j*+1)_ for all neurons in layer *j* + 1, are increased by an amount

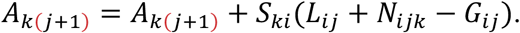

In order to balance total excitation and inhibition from layer to layer, the following summation equation, justified by the physical requirement of stability in the system, is enforced

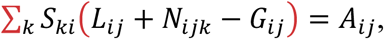

which ensures excitation net of inhibition received by the firing neuron is neither amplified nor attenuated as it is transmitted to the next layer. In order to control the relative strength of the non-linear NMDAR-mediated conductance to the linear terms in the model, we introduce the modeling parameter *p* and enforce

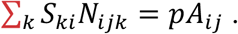

Since, by definition, ∑*_k_ S_ki_* = 1, we have that *L_ij_* − *G_ij_* = (1 − *p*)*A_ij_* and it follows from the previous two equations that the ratio of non-linear conductance to linear conductance net of inhibition will be precisely *p*/(1 − *p*).

In order to model the opening and closing of the NMDAR gates with the post-synaptic firing average, we invoke a logistic function which smoothly and monotonically transitions from a value of 0 (gates fully closed) to a value of 1 (gates fully open),

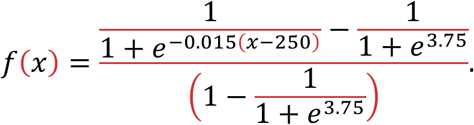

We can then determine the non-linear conductance, *N_ijk_*, as

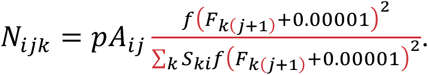

Where *p* is the percentage of non-linear component in the post-synapses, the function *f* is a monotonically increasing form of a logistic function given by

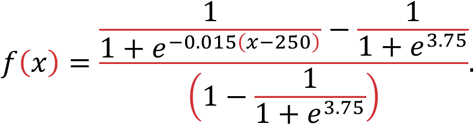

The value 0.00001 is merely chosen as a small value that will generally have insignificant impact on the calculation but prevents the denominator from being 0 before any neurons have fired in the simulation.

Following these definitions, the three postsynaptic components, corresponding to AMPAR- and NMDAR-mediated excitation and GABAAR-mediated inhibition, respectively, together sum over layer *j* + 1 to be precisely the value of *A_ij_* which ensures the stability of the algorithm or excitation/inhibition balance. If they summed, in general, to be greater than *A_ij_*, then in each subsequent layer the number of times the neurons will fire grows exponentially and unphysically. If they summed, in general, to be less than *A_ij_*, then in each subsequent layer the number of times the neurons will fire shrinks exponentially and neurons may almost never fire in the furthest layers, which is also unphysical. After the values of *A_k_*_(*j*+1)_ have been updated, the value of *A_ij_* is set to zero and the algorithm moves on to the next neuron to fire. For neurons firing in the last layer, there is no transmission, and when neuron *i* in the last layer fires, we need only to set *A_i_*_30_ to zero. The linear (AMPAR-mediated) conductance does not depend on how active any neuron is, while the non-linear (NMDAR-mediated) conductance increases as the receiving neuron becomes more active relative to the other connected neurons. Even though the non-linear conductance is only 27.5% (*p* = 0.275) of the total, this is sufficient for neuronal priming to be observed.

### Distribution of c-Fos expression level in mice that were subjected to trace fear conditioning

Control mice, including vehicle control and genotype control at nine weeks after tamoxifen/vehicle administration, were euthanized 45 minutes after trace fear conditioning following the protocol above. Sliced brain samples (50 μm) were subjected to fluorescence immunostaining using primary antibodies against c-Fos (1:5000, Cat# RPCA-c-Fos, EnCor) and secondary antibodies (conjugated with Alexa Fluor 488, 1:500) to measure neuronal activity level in the hippocampal CA1 region. All fluorescence images were captured by a Nikon A1R HD confocal microscope and acquired using the same settings including exposure time, gain and laser power settings. The fluorescent intensity of c-Fos signal from individual cell was measured by Fiji ImageJ - Analysis Particles ^57^. The c-Fos fluorescent intensity of the individual neurons was normalized to the average of all c-Fos positive neurons in the same images in order to plot the histogram of c-Fos activity level responding to trace fear conditioning.

### Pharmacological studies using ketamine in conjunction with *in vivo* hippocampal calcium imaging and trace fear conditioning

To examine the roles of NMDAR in controlling neuronal priming, we used ketamine to block NMDA receptors and examined how receptor blockade affects neural activity hierarchy. Two different timings were chosen to administer ketamine (10 mg/kg) to observe various effects of ketamine: (1) 5 min before conditioning. Afterwards, mice were subjected to one round of trace fear conditioning (7 cycles), and they were still under the influence of ketamine during the training. This was followed by a recall testing 2-3 hours later. This experiment was to observe how ketamine affects associative learning and hippocampal neural dynamics. (2) Mice were subjected to two rounds of trace fear conditioning (7 cycles + 7 cycles). Ketamine was injected ∼5 minutes after the 1^st^ round of training. Afterwards, mice were put back to home cages for rest and recovery. There are clear differences in behaviors when animals are under ketamine control or not. 2-3 hours later when ketamine effect faded away ^39,40^ and mice stayed fully awake and behaved normally, they were subjected to the 2^nd^ round of training. The work of ketamine injection between two-round of training was to observe if ketamine affects the existing neural hierarchy that manifested in the 1^st^ training. The 2^nd^ training was implemented to identify whether primed neurons ensemble was reshuffled by ketamine.

### Statistical analysis and graphic data presentation

If the data’s sample sizes were more than 50, or if the sample sizes were less than 50 but they passed normality tests, statistical analyses were carried out using Student’s *t*-test for a two-group comparison, or ANOVA test for multiple-group comparison. If the data’s sample sizes were less than 50 and they failed to pass normality tests, a nonparametric test was used for two-group comparison or for multiple-group comparison. In figures, ns indicates statistically not significant, * *P* < .05, ** *P* < .01, *** *P* < .001, and \*\*\*\**P* < .0001. A difference in the data comparison was considered statistically significant if *P* < .05. In the graphics, data points are shown as the means ± the standard error of the mean. Results were analyzed and presented by GraphPad Prism 9, home-made MATLAB codes, Clampfit 10.3, CorelDraw, and Adobe Premiere Pro.

## Results

### Instant one-day fiber-optic *in vivo* calcium imaging avoids pathological neuronal activation, uniquely suited to determine neural activity hierarchy

It is critical to ensure there is no pathological neuronal activation before the *in vivo* calcium imaging, as this helps maintain neural activity at the physiological state to allow for the measurement of the neural activity hierarchy and the quantitative identification of primed neurons. We combined a deep-brain fiber-optic confocal (FOC) fluorescence endomicroscopy with mouse trace fear conditioning experiment in a time-locked manner. One limitation of this imaging system is that although it can maintain imaging stability for more than 7 hours, it does not permit repetitive imaging for multiple days ^20^. However, this instant one-day imaging approach instead confers one advantage: it omits the need for GRIN lens implantation into the brain, thereby avoiding pathological activation and hippocampal inflammation caused by GRIN lens implantation and tissue injury (Fig. 1A). Indeed, the mouse’s hippocampus did not display pathological activation weeks after canula installation surgery and AAV injection, as evidenced by comparable c-Fos expression and GFAP staining in the surgical and non-surgical hemispheres (Fig. 1B-E). In contrast, 2-3 days after completion of imaging, GFAP expression was found to be strongly elevated in the imaged region, compared to the non-imaged hemisphere (Fig. 1F). This demonstrates that the mouse’s hippocampus stayed very close to the physiological conditions right before imaging, but not after the imaging that could cause certain injury.

**Fig. 1.**
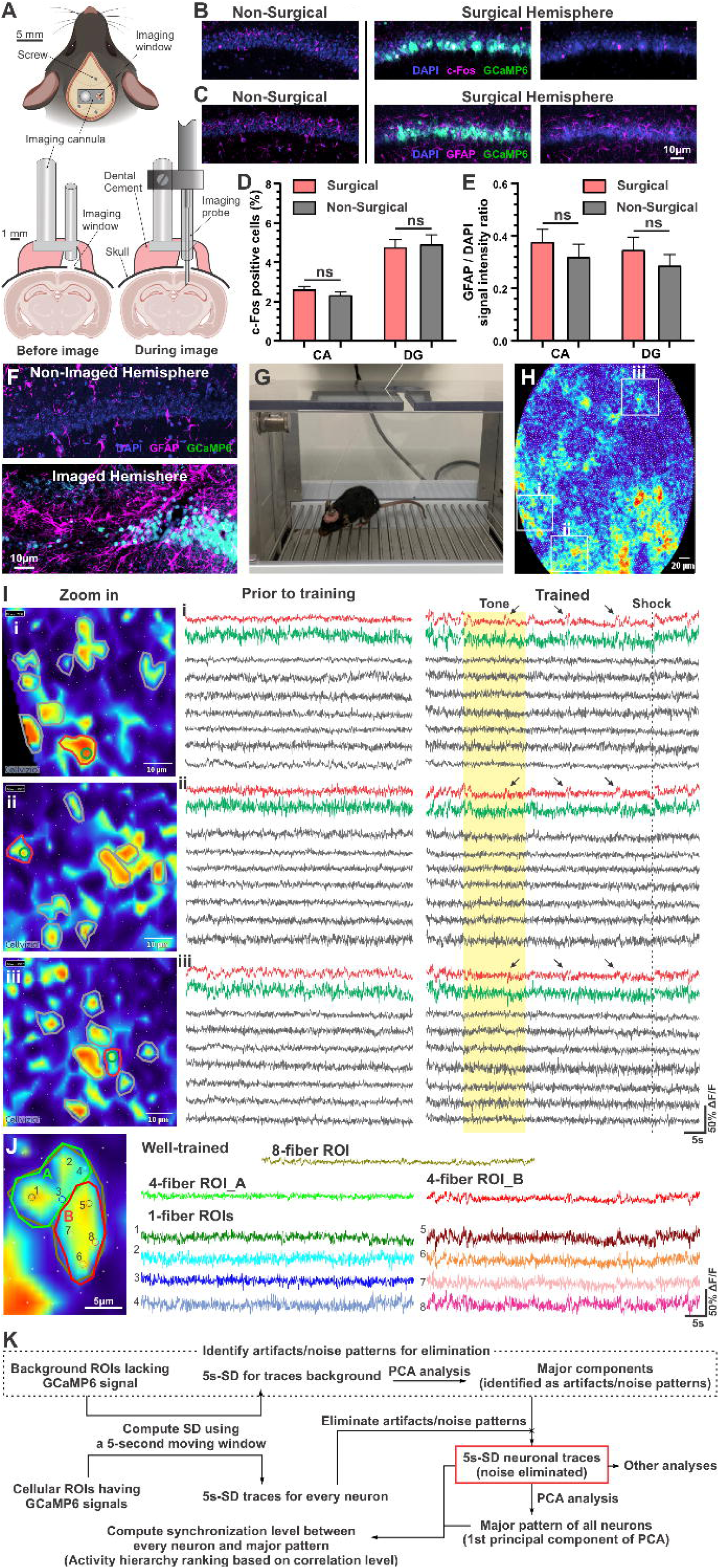
Minimally invasive surgery and one-day imaging at the cellular resolution avoids hippocampal inflammation and pathological neuronal activation. (A) Top: Illustration of the surgical setup where the imaging window aligned with an imaging cannula, which were stabilized by screws and dental cement. Bottom: A cross-section of the setup before and during imaging. Cannula implantation surgery was installed above the skull and surgery only slightly affected the surface of neocortex, but not the hippocampus. A tiny needle-like imaging microprobe (∅ = 0.47 mm) reached the dorsal hippocampus through the imaging cannula. Surgery pictures were created using BioRender (www.BioRender.com). (B) Immunofluorescence staining using c-Fos antibody on hippocampal slices 8 weeks after AAV-Syn-GCaMP6m injection and canula stereotaxic surgery and right before *in vivo* fiber-optic imaging. Left: non-surgical hemisphere showing c-Fos positive neurons. Right: surgical side from same brain slice with GCaMP6m expression (green), showing similar levels of c-Fos expression (magenta). (C) GFAP staining (an indicator for reactive astrogliosis) shows no differences between the surgical and non-surgical hemispheres. (D-E) Statistics of c-Fos-positive neuron number (D) and GFAP fluorescence intensity (E) relative to DAPI in non-surgical and surgical hemispheres. There was no difference between non-surgical and surgical hemispheres, paired Student T-test. (F) Three days after imaging, imaged hippocampal tissues (right) displayed signs of astrogliosis, as evidenced by heightened expression of GFAP. The left non-imaged hemisphere had no signs of gliosis. (G) A mouse carrying an imaging probe could behave freely. (H) A whole view of endoscopic image. 3 distantly located areas (white boxes) were selected and enlarged in (I) to demonstrate the single cell resolution of imaging. (I) In each area, one primed neuron (red ROI and red trace) was surrounded by several non-primed neurons (grey ROIs and grey traces). Green traces were calcium signals collected from one fiber ROIs located at the center of red ROIs. One-fiber signals (green ROIs and green traces) had a lower signal-to-noise ratio, but their dynamics were very similar to multiple fibers signals (red traces) that covered a larger area. Primed neurons were randomly active prior to training but had bursting when the mouse was trained to engage in conditioning. These separately distributed primed neurons (red) also synchronized with each other (i, ii and iii), whereas neighboring non-primed neurons (grey) showed little bursting patterns or synchronization after being trained. (J) An empirical method to identify single cell ROIs. An 8-fiber ROI covering a large area and more than one neuron (golden). Individual fibers are shown in the left image. Calcium traces of each fiber (#1-8) are shown in the right, which can be classified into two groups (A and B). Neuron A encompasses fiber 1-4 (all showing similar calcium dynamics), while neuron B contains fiber 5-8 (all showing another pattern). Thus, the 8-fiber ROI was interpreted to cover 2 neurons (A and B). (K) A scheme showing the procedure of imaging data pre-processing and subsequent data analysis.

The imaging microprobe has an inner diameter of 300 µm, encasing a bundle of ∼7600 optic fibers of 1 µm. It has a lateral resolution of 3 µm, which allowed for the tracking calcium dynamics at a cellular resolution ^20^. We also developed multiple empirical strategies to ensure that our imaging data was collected from single cells (Fig. 1H-J). ROIs encircling multiple fibers (minimal 3 fibers) (outer part, red) or 1 fiber (inner part, green) were manually drawn respectively, and their calcium traces were compared. 1-fiber ROIs covered a smaller, central part of the soma, whereas multiple-fiber ROIs covered a larger area (∼ 7 x 10 µm) (Fig. 1I). The fluorescence intensity at the center is brighter than the edge, which helped separate two neurons. We also compared active neurons with several relatively inactive neurons surrounding it. During training, neuronal synchronization occurs among active neurons which generally locate distantly, and rarely among neighboring neurons (Fig. 1I). To distinguish calcium signal from possible interference by motion artifacts, we filtered imaging data with a 1 Hz high-pass Fourier filter. The synchronized bursts were clearly retained in filtered traces (Fig. S1). We mostly used multiple-fiber ROIs data in our subsequent analysis because this yielded a better signal-to-noise ratio (Fig. 1I). However, multiple-fiber ROIs may potentially cover a region having two neurons. To address this, if a multiple-fiber ROI’s activity pattern was very similar to all individual 1-fiber ROIs within the area, we deduced that the multiple-fiber ROIs cover a region from the same neuron as those 1-fiber ROIs (Fig. 1J). If the activity pattern of multiple-fiber ROI was different from individual 1-fiber ROIs, we reasoned this multiple-fiber ROI may cover a region for more than one neuron (Fig. 1J), whose data were then excluded in our analysis, or the ROI could be re-drawn until a single neuron was clearly defined. Further, we excluded imaging regions if two neurons’ imaging intensity contrast could not be clearly separated. Hence, we were able to collect imaging data from ROIs of single neurons and exclude ROIs that cover more than one neuron. Together, this deep-brain imaging at the cellular resolution in freely behaving mice free of prior pathological activation is well-suited to measure hippocampal neural activity hierarchy *in vivo*.

Fluorescent traces of cellular calcium signals were collected and transformed into 5 second moving standard deviation (SD) traces, followed by artifact/noise elimination, prior to computing synchronization levels and performing other statistical analysis (Fig. 1K). More specifically, we first converted calcium traces into 5-second moving SD traces, if not otherwise specifically indicated, to smooth white noise and amplify bursts. Next, we calculated the projection of the individual neuronal SD traces onto the identified background noise patterns, which result from PCA to the SD traces of 12 background ROIs, and subtracted that projection from individual neuron’s SD traces, thus orthogonalizing individual neuron’s SD traces to all background patterns (see Methods). After eliminating background patterns, the SD traces of individual neurons were used for the subsequent statistical analysis, including extracting synchronized patterns among all neurons (or among selected groups) to generate the major pattern of the PCA, to compute the correlation coefficients of individual neurons with the major pattern, to calculate the correlation between mouse freezing behaviors and the major pattern, and to compute the SD-of-SD that reflects neuronal activity levels, as well as to identify primed neurons using synchronization levels and the SD-of SD values. ROI data also contain the region’s X-Y coordinates information that was used to plot the relative location of individual neurons under an imaging field.

### Burst synchronization is a key feature of trace fear conditioning

To help verify our methods to determine neural activity hierarchy and identify primed neurons, we used Ift88 cilia KO mice (Ift88 flox/flox; UBC-Cre/ERT2 mice, tamoxifen-induced), which represent a temporally induced primary cilia loss-of function animal model ^53,58^. Ift88 cilia KO mice have severe learning deficits in Morris water maze test and trace fear conditioning test ^55^. We subjected control and cilia KO mice to trace fear conditioning, conducted in conjunction with *in vivo* calcium imaging. We monitored the calcium dynamics of hippocampal neurons during the whole conditioning procedure including training and recall testing, prior to training and prior to recall, as well as under isoflurane-induced anesthesia for imaging normalization. Prior to training, both control’s and cilia KO’s calcium traces exhibit irregular dynamics (Fig. 2A-B). At training cycle 1 (cy1), calcium traces of the control animal started to show a few bursts, but the correlation among neurons was very weak. Starting from the 4^th^ training cycle, the animal behavior shows a strong difference between control and KO mice. Calcium traces from the control animals formed obvious synchronization. A strong burst synchronization correlating with animal behavior was formed among control neurons at the 7^th^ training cycle. After resting for 2 hours, these neurons went back to randomly active, similar to that prior to training. Upon recall, they underwent tone-induced synchronization quickly (Fig. 2A). In contrast, the cilia KO mouse does not form a clear synchronization during training or recall (Fig. 2B). We also constructed frequency density plots using these calcium traces after eliminating background and motion artifacts to highlight the high-frequency signals. Consistently, control neurons exhibited bursting patterns in training cy4 - cy7 as well as during recall testing, whereas cilia KO neurons failed to burst (Fig. 2A-B bottom). F-test confirmed there were significant differences in variability during training cy4 - cy7 between control’s and cilia KO’s dataset (not shown). This data indicates that burst synchronization of high-activity neurons is a key feature of trace fear conditioning.

**Fig. 2.**
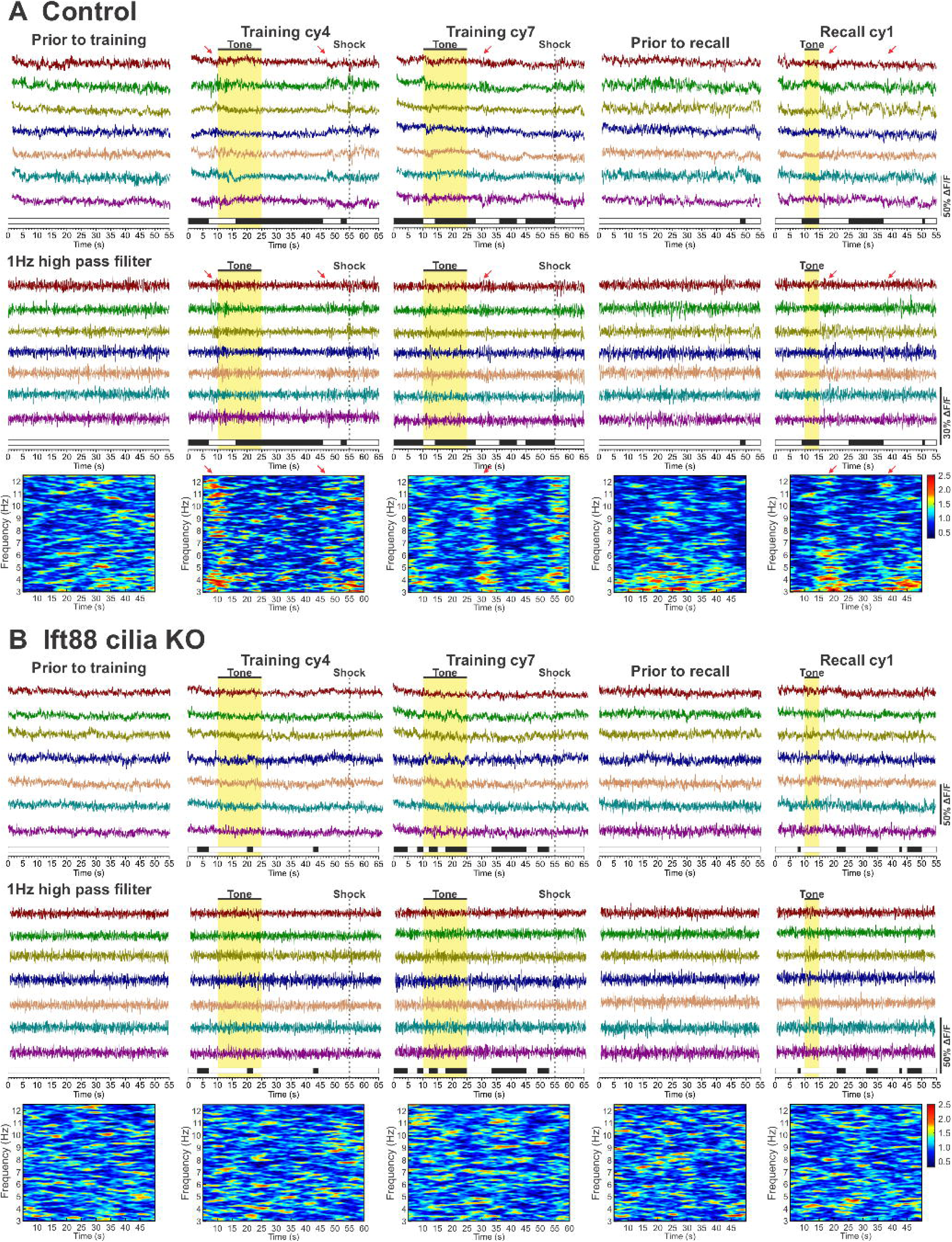
High-activity hippocampal neurons of control mice, not Ift88 cilia KO mice, develop trace memory-associated burst synchronization. **(A) (Top)** Representative calcium raw traces of 7 high-activity neurons from a control mouse. (Middle) Calcium imaging traces were filtered by a 1 Hz high-pass Fourier filter. Time bars: mouse moving (white), freezing (black). (Bottom) Time-frequency density plots of calcium dynamics of 7 high-activity neurons. **(B)** (Top) Representative calcium raw traces of 7 neurons with high activity from one cilia KO mouse. (Middle) Calcium imaging traces were filtered by a 1 Hz high-pass Fourier filter. (Bottom) Time-frequency density plots of calcium dynamics. High-activity neurons of control mice developed burst synchronization in late training cycles and upon a successful recall, whereas those of cilia KO mice don’t display bursts or any specific pattern.

### Identifying primed neurons based on neurons’ correlation levels with the major pattern of PCA

We previously observed that repetitive trace fear conditioning modified the calcium dynamics of high-activity neurons from irregularity to burst synchronization (Fig. S3A). This also resulted in changes of the correlation level distribution. After eliminating background pattern from individual neurons, we extracted a major pattern of PCA of all neurons. We calculated Pearson correlation coefficients of every neuron with the major pattern (Fig. 3A). In control mice, the correlation coefficient of all neurons displayed a relatively linear distribution prior to training, meaning these neurons were evenly distributed on the 1^st^ principal component. After being trained, high-activity neurons synchronized with each other to dominate the major pattern and increased their correlation with the major pattern of PCA. In contrast, cilia KOs did not exhibit marked changes over training and recall testing, and the correlation coefficients of all neurons remained relatively unchanged (Fig. 3A). Fig. 3A also demonstrates that trace fear conditioning remarkedly increased the percentage of neurons with correlation levels higher than 0.7 in control mice, but not in KO mice.

**Fig. 3.**
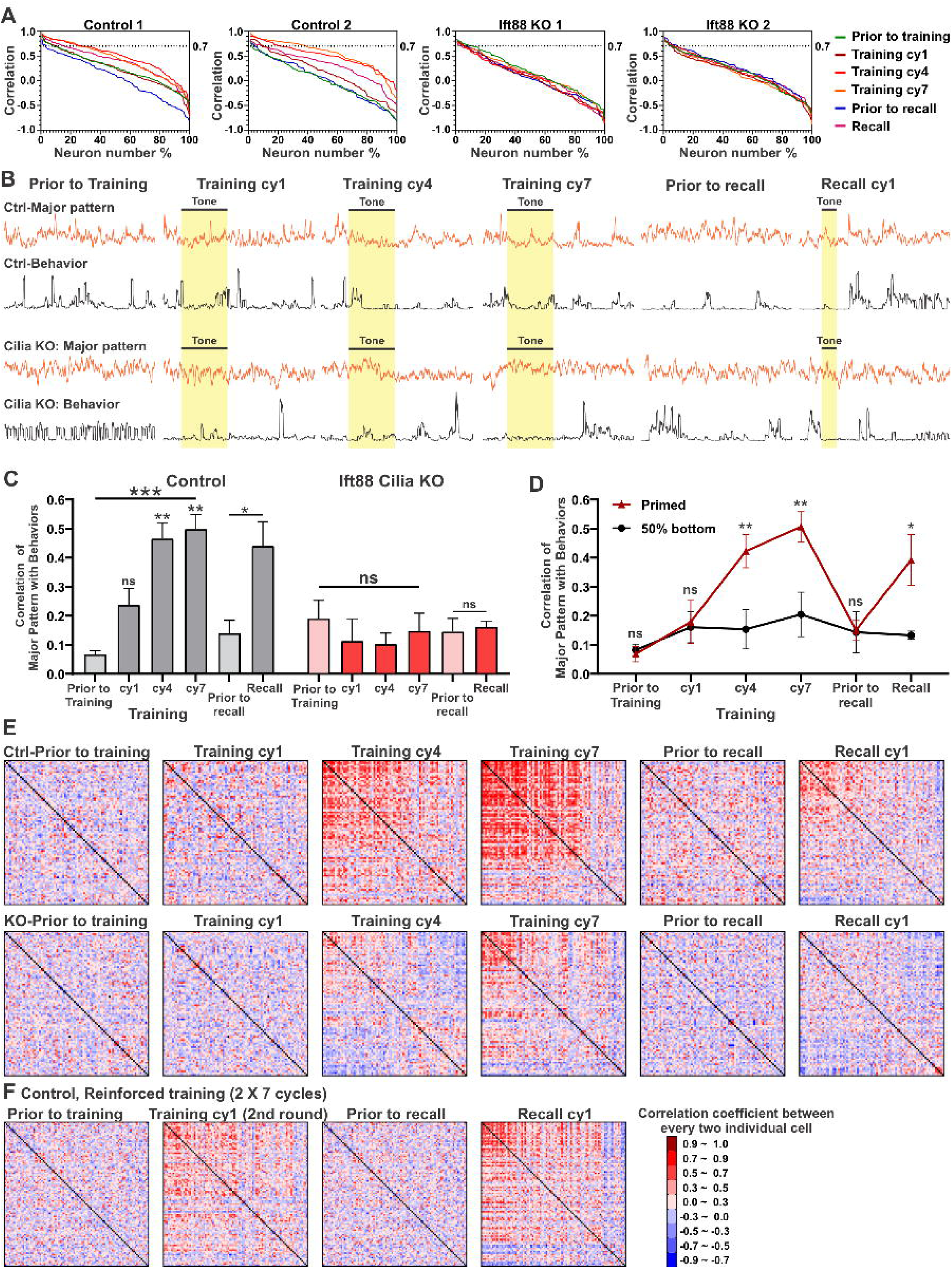
Synchronization of primed neurons is correlated with animal conditioned freezing behavior in control mice, but not in Ift88 cilia KO mice. **(A)** Pearson correlation coefficient of individual neurons compared to their major patterns of PCA at different conditions. Examples from two controls and two cilia KO mice are shown. Correlation levels with the PCA major pattern increases with training and upon recall testing in control animals, but not in KO animals. **(B)** Top: Major patterns (orange) computed from 0.5s-moving SD trace of all neurons and freezing behavior (black) of one control mouse. The control animal had a clear response to tone and shock when trained. Moreover, the major pattern showed a clear response to stimulus and a strong correlation with mouse behavior. Bottom: major pattern of 0.5s-moving SD trace of all neurons and freezing behavior of one cilia KO mouse. The major pattern of this cilia KO animal cannot develop an obvious burst. **(C)** Pearson correlation coefficient between the major pattern of all neurons and behavior data (n = 5 pairs, mixed sexes). The correlation levels were increased with training and upon recall in control animals, but not in cilia KO animals. Repeated measure one-way ANOVA with post hoc Tukey’s multiple comparisons test between prior to training and each other cycle. Paired Student T-test between prior to recall and successful recall. **(D)** Pearson correlation coefficient of primed neurons’ and bottom 50% non-primed neurons’ major patterns with behavior data in control animals (n = 5, mixed sexes). The correlation levels of primed neurons were significantly higher than the bottom 50% neurons during training and upon recall. Bonferroni’s multiple comparisons test of two-way ANOVA, primed neurons vs bottom 50% neurons. ns, not significant; *, p<0.05; **, p<0.01; ***, p<0.001. **(E)** Heatmap of correlation levels between every two neurons from a control mouse (top) and a KO mouse (bottom). The same neuron was placed at the same position in these heatmaps. In the control mouse, the same group of neurons displays high correlation levels during training and recall testing, whereas neurons of the cilia KO mouse did not form strong synchronization with training. **(F)** Enhanced training promotes neuronal synchronization during recall in controls. Reinforced trace fear conditioning paradigm: 7 cycle training (not imaged) - 2 hours rest - 7 cycle training again (imaged) - 3 hours rest - recall (imaged). Pairwise neuron correlation levels are indicated warm color.

We classified all neurons based on their Pearson correlation coefficients compared to the major pattern of all neurons. As a rule of thumb, the correlation levels higher than 0.9 were defined as very highly correlated; the correlation levels between 0.7 to 0.9 were counted as highly correlated; the correlation levels between 0.5 to 0.7 were counted as moderately correlated ^56^. We observed that high-activity neurons started to show burst synchronization in the middle of training (e.g., cy4) and can last until the end of training (cy7). In recall, tone-induced synchronization is also crucial for memory retrieval. Thus, we summarized the correlation levels in the middle of training (i.e., the 4^th^ training cycle), after well-trained (generally the 6^th^ or 7^th^ training cycle) and a successful recall (whichever 1^st^ or 2^nd^ recall testing) of every neuron to be the level for identification. The cut-off was set as follow: individual neurons with sum of correlation coefficients higher than 2.0 was defined as primed neurons; others were non-primed neurons, including silent neurons (sum of correlation level between -1.5 to 1.5) and intermediately active neurons (sum of correlation level between 1.5 to 2.0). After the classification, we found that primed neurons dominated the major patterns extracted by PCA when mice were actively engaged in the associative learning and memory recall testing (Fig. S4). In contrast, KO mice did not have many primed neurons, which did not dominate the major pattern associated with trace fear conditioning (Fig. S4).

### Synchronized activity of primed neurons highly correlates with the freezing behaviors of control mice, but not cilia KOs

To examine how hippocampal neurons collectively engage in trace fear conditioning, we also ran a PCA using time-windowed SD trace of all neurons collected from control and cilia KO mice to extract the major patterns of every cycle. These patterns revealed the major bursting responses of all neurons, which were dominated by the group of primed neurons, as shown in Fig. S4. We compared the correlation level between the major patterns with mouse freezing behaviors. Behavior results were also displayed as moving SD traces, following the same time-window as calcium traces. The representative pattern of moving SD calcium traces and mouse behaviors of a control animal exhibited strong coherence responding to tone during trace fear conditioning and recall (Fig. 3B Top), whereas cilia KO neurons did not (Fig. 3B Bottom). We quantified the correlation levels of mouse behaviors with the corresponding major patterns of control and cilia KO mice. Prior to training, both control and cilia KO mice had only a weak correlation between behaviors and the major pattern. Control mice exhibited increasing correlation levels during training or recall testing, whereas cilia KOs had little changes over the training and recall (Fig. 3C).

If our classification of primed neurons is sound, then identified primed neurons should dominate the correlation with mouse freezing behavior. To verify this, we employed PCA to extract the major pattern selectively from primed neurons’ signals. We also extracted the major pattern using SD traces from the bottom 50% non-primed neurons. We conducted a correlation analysis between major patterns of primed and non-primed neurons with mouse locomotion data, respectively. Fig. 3D demonstrates that the correlation between primed neurons’ major pattern with animal locomotion markedly increased with training, then returned to the basal level during rest, and increased again upon recall testing. In contrast, the major pattern of non-primed neurons did not show such a correlation pattern with behaviors during training or upon recall. It is worth mentioning that we were unable to identify enough primed neurons from each cilia KO mouse to permit PCA to extract any pattern. Taken together, these data confirm that the synchronized activity of primed neurons highly correlates with the freezing behaviors during the training and recall testing in control mice.

To examine if neural synchronization is maintained among the same group of neurons, we built heatmaps using pair-wise neuronal correlations from training to recall testing. All neurons were ranked following the sum of their Pearson correlation coefficient with the major pattern (middle of training, i.e., cy4, end of training, i.e., cy7, and a successful recall), the same factor used for primed neuron classification.

These neurons were kept in the same position in the heatmaps from training to recall, and neurons having a higher sum of correlation coefficients were put to the top left. In the control mouse, neurons in the top-left corner were highly synchronized with each other. Pair-wise neuronal synchronization started to take shape in training cy4 and formed a stronger correlation when the mouse was well-trained in cy7. Moreover, the pair-wise neuronal synchronization partially re-emerged during recall testing (Fig. 3E-top). In contrast, cilia KO neurons had much weaker changes in pair-wise neuronal synchronization during training and recall testing (Fig. 3E-bottom). This data suggests that neuronal synchronization correlates with mouse freezing behaviors of control mice, which is defective in cilia KOs. One concern is that after rest, the neuronal synchronization during recall is not very strong. To enhance memory formation and achieve a stronger neuronal synchronization during recall testing, a reinforced trace fear conditioning was applied to control animals. This group of animals were subject to two rounds of trace fear conditioning prior to recall testing. Indeed, this enhanced training markedly increased the pair-wise neuronal correlation during recall testing (Fig. 3F). Together, these results support the conclusion that neuronal synchronization is well maintained among primed neurons in control mice, which is closely associated with trace fear memory formation and retrieval.

### Ift88 cilia KO mice display drastically reduced hippocampal neuronal activity

We converted neuronal calcium traces into 5-second moving SD traces, which reflect the magnitude of the fluctuations of calcium dynamic. A trace with a bursting pattern has significant variation in the SD trace over time. The variation in the SD trace in time can be measured by taking another standard deviation of the entire SD trace (Fig. S2A). This new quantity, which we named SD-of-SD, gives an indication of the variation in the activity level over the course of the experiment (Fig. S2B). We noted that some high-activity neurons tend to form a burst when responding to stimulus or spontaneously when a mouse is trained to engage in trace fear memory ^20^ (Fig. S3). The SD-of-SD, as a measure of “bursting” can be used as alternative method to approximate neuronal priming. For instance, a neuron that exhibits “bursting”, forming a peak when responding to learning cues or spontaneously occurring, followed by a period of low activity, have a high SD-of-SD, while neurons that are either consistently active, consistently inactive, or simply white noise at any amplitude have low SD-of-SD (Fig. S2B). Since the computation of neuronal activity levels was changed from variance to SD-of-SD, the apparent distinctions between primed neurons and non-primed neurons required a heavier skewed tail, which switched fitting functions from log-normal probability density function to log-logistic probability density function ^59^.

We plotted the histograms of the activity levels of individual neurons measured by the SD-of-SD under different conditions (Fig. 4A). Similar to our previous report ^20^, the neuronal activity histograms are clearly right skewed. Based on the histograms, most neurons having low activity levels were classified as non-primed neurons, whereas a small portion of neurons with SD-of-SD ratio higher than 3 were considered candidates for primed neurons (Fig. 4A). In control mice, the fitting curves were significantly shifted to right by training, suggesting an increase in bursting activity (Fig. 4A-i). It is worth noting that in the right-skewed tail, the number of high-activity neurons was dramatically increased, because of neuronal bursting caused by training. Similar to training, recall testing also led to a clear rightward shift in the neuronal activity levels in control mice (Fig. 4A-ii).

**Fig. 4.**
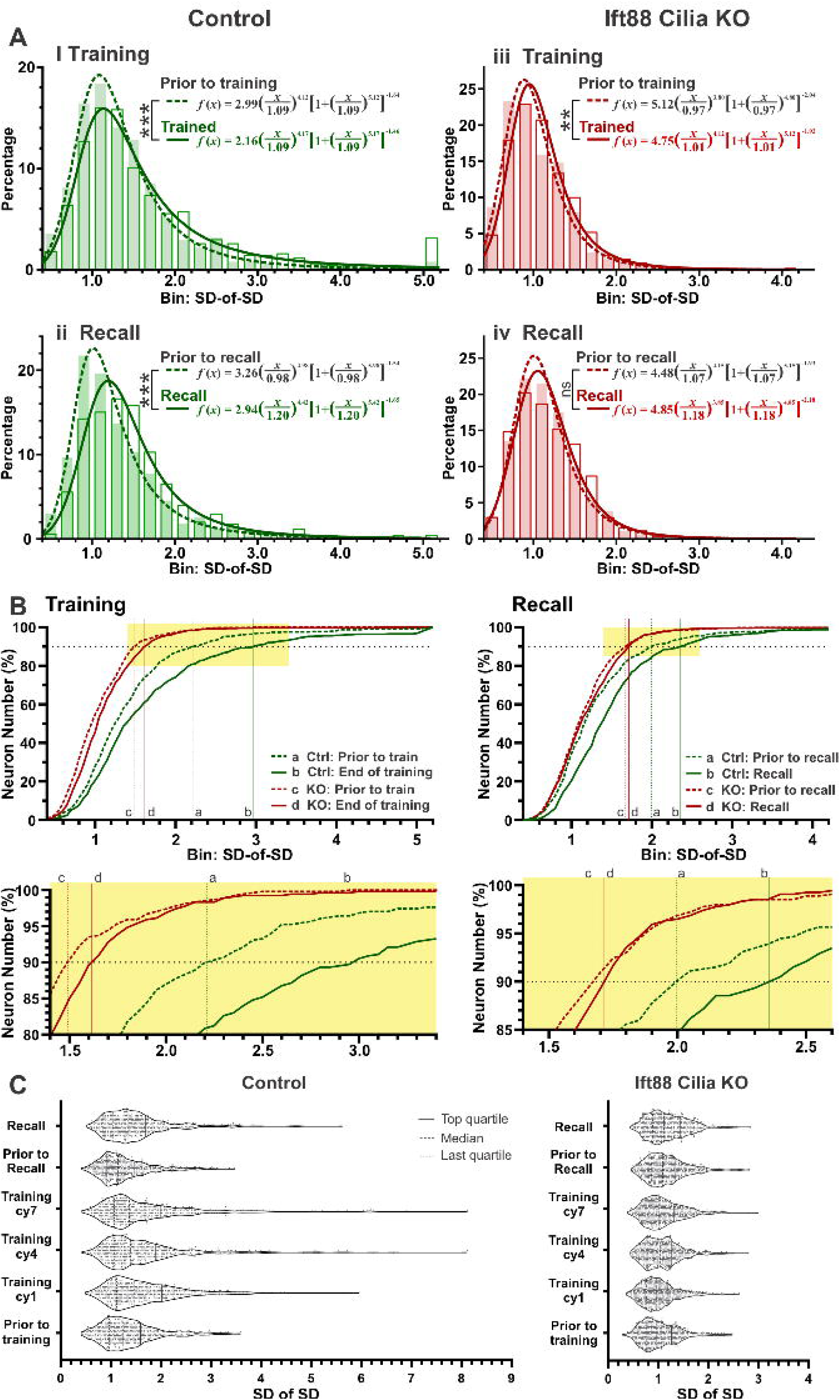
Reduced hippocampal neuronal activity in Ift88 cilia KO mice. **(A)** Activity histograms of individual neurons in control mice (left, green) and cilia KO mice (right, red). The SD-of-SD of individual neurons prior to training (top, dashed), at training cycle 7 (cy7) (top, solid), prior to recall (bottom, dashed), and during recall (bottom, solid); all is normalized to those under anesthesia. The fitting curves were made by probability density functions; K-S tests were used to compare the differences. Controls, n = 506 neurons from 5 control animals. K-S test between Prior to training and the end of training, ***, p < 0.001; prior to recall vs successful recall, ***, p < 0.001. Cilia KOs, n = 541 neurons. K-S test between prior to training and the end of training, **, p = 0.004; prior to recall vs successful recall, ns, p = 0.13. **(B)** Cumulative distribution of the SD-of-SD of individual neurons in controls (green) and cilia KOs (red). The guides were set on y = 90% to highlight the neurons having the top 10% of SD-of-SD levels. The extended lines labeled the intersections between every curve and the guides. The highlighted boxes were zoomed in at the bottom. **(C)** Violin plots of SD-of-SD of calcium dynamics of individual neurons at different conditions. Left, controls; Right, cilia KOs. Data were collected from 506 neurons out of 5 control animals and 541 neurons out of 5 cilia KO animals. The SD-of-SD shows a rightward shift with training or upon recall in control animals, but not in cilia KOs.

We used cilia KO mice as a negative reference to assess our estimation of primed neurons. The neuronal activity histograms (measured by SD-of-SD) of cilia KOs were weakly right-skewed, indicating a reduced number of primed neurons in cilia KOs. Moreover, trace fear conditioning still caused a rightward shift in the histogram in cilia KOs, but not as strong as controls (Fig. 4A-iii). This data was consistent with impaired memory behaviors (Fig. 2). In recall, neurons in KO mice showed little rightward shift (Fig. 4A-iv). To further compare the activity levels of high-activity neurons that fall into the right-skewed tail, we calculated the cumulative distribution for the SD-of-SD and focused on the top 10% active neurons (Fig. 4B). The extended line from the intersection of each curve and the subline for y = 90% revealed the activity level of top 10% neurons in each group. In control mice, the cut-off for top 10% highly active neurons was increased from 2.22 (prior to training) to 2.98 (when trained) in trace fear conditioning. In cilia KOs, this reduced increase was from 1.49 (prior to training) to 1.63 (trained). Prior to recall, the top 10% cut-off for control mice was 2.0 and increased to 2.38 by the recall cycle. However, cilia KOs exhibited a weaker increase from 1.68 to 1.71 during recall. Additionally, the violin plots of SD-of-SD further confirm a similar result that the neuronal activity levels of control animals shifted dramatically by training and recall, whereas cilia KO neurons had little change (Fig. 4C).

Since both high neuronal activity levels and memory-associated burst synchronization are key features of primed neurons, we combined these two factors and plotted SD-of-SD against correlation coefficients (with the major pattern) to verify the identification of primed neurons. In Fig. 6A, the cut-offs were set as SD-of-SD higher than 3 and the correlation coefficient higher than 0.7. In control animals, a cluster of primed neurons were identified as they appeared in the upper-right corner in training cy4 to cy7 and re-appeared in recall testing. In contrast, cilia KO animals failed to find many primed neurons in the upper-right corner throughout the training and recall cycles, demonstrating significantly lower number of primed neurons. We further mapped out the *in-situ* positions of identified primed neurons (Fig. 5B-D), along with silent neurons, on imaging fields. Primed neurons are found to sparsely distributed throughout imaging fields (Fig. 5B-C), and they had an average neighboring distance of ∼40 µm (Fig. 5E). This study confirms that under an *in vivo* imaging probe, memory-eligible primed neurons do not cluster together.

**Fig. 5.**
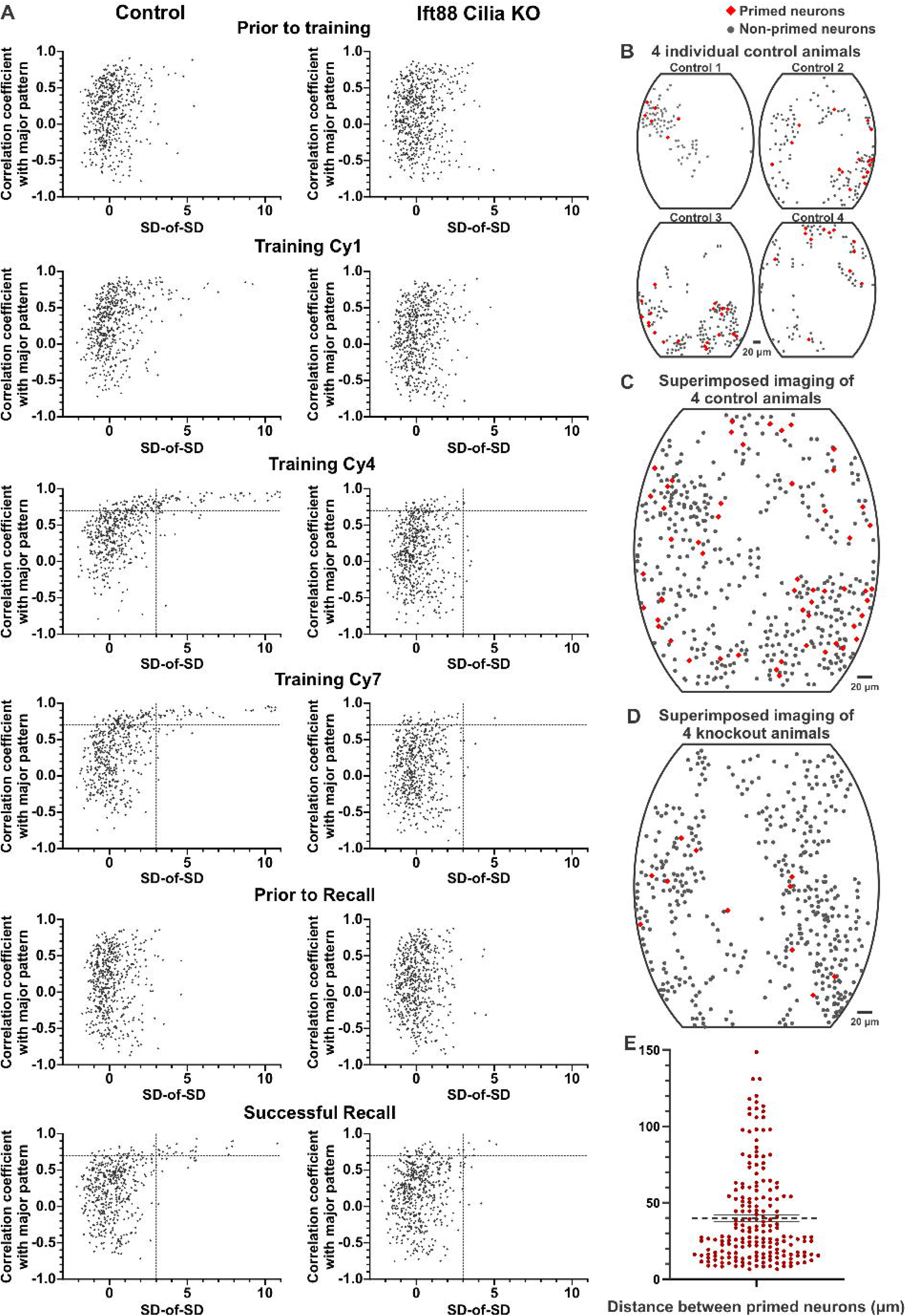
Plotting neuronal synchronization against SD-of-SD to identify primed neurons. **(A)** Pearson correlation coefficient (to the major pattern) vs the activity level (SD-of SD) of individual neurons. Controls (left) and cilia KO mice (right). X-axis represents the SD-of-SD of individual neurons normalized to the basal level of isoflurane-induced anesthesia, Z-scored. Y-axis is the correlation level of individual neurons to the major pattern of PCA. Dashed lines at Y = 0.7: neurons with synchronization levels higher than 0.7 were considered high synchronization; Dashed lines at X = 3: neurons having a SD-of-SD higher than 3 were viewed as high activity. Neurons in the top-right (with high correlation with the major pattern and high activity) were defined as primed neurons. Based on the combined calculation from cy4, cy7, and a successful recall, control mice have markedly more primed neurons than cilia KO. **(B-D)** Sparse distribution of primed neurons *in situ*. Superimposing four individual images **(B)** into one imaging field, controls **(C)**, cilia KOs **(D)**. Individual cells were scattered over the imaging-field and identified primed neurons were sparsely distributed throughout the imaging-field. **(E)** Distance between every primed neuron with its three closest primed neurons in control animals.

**Fig. 6.**
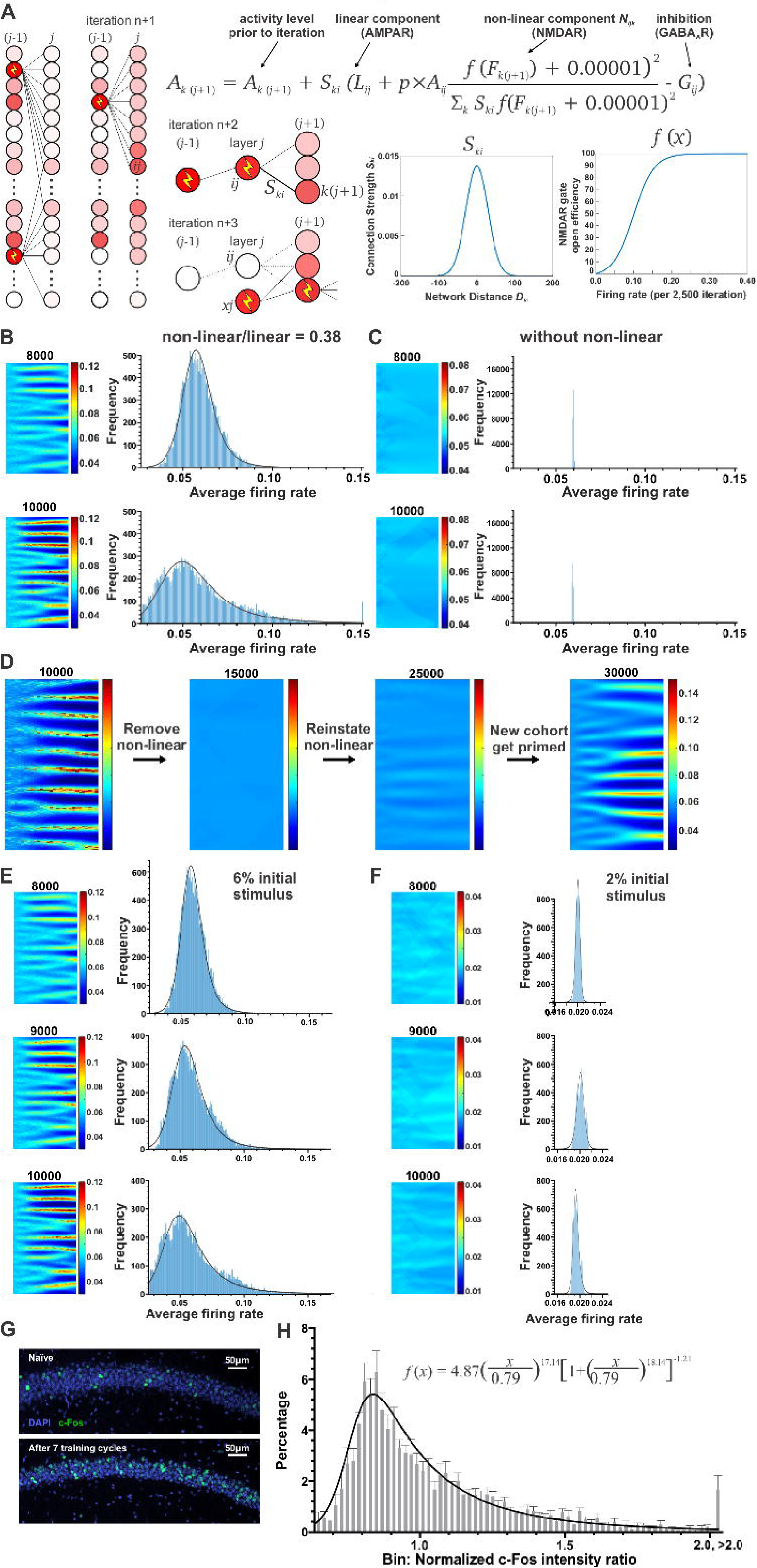
Accumulation of non-linear weighted synaptic transmission naturally leads to activity hierarchy in a simulated neural network. **(A)** Neural network simulation. A fired pre-connected neuron (*Nij* at row i, column j) releases neuronal transmitters from pre-synapse, the post-synaptic conductance of the next layer neuron (*Nk*(*j*+1)) is mediated by AMPAR (the linear component) and NMDAR (the nonlinear component) and GABAAR (the major inhibitory component). The connection strength *Ski* is dependent on the distance between neurons. The transmission efficiency of the non-linear conductance is dependent on the opening rate of the NMDAR channel, which are affected by post-synaptic neuronal activity. **(B)** Initial neuronal activity was set to 6%. Firing rate for the entire simulation during 8,000 and 10,000 computation iterations (left), showing pattern development and neural priming in the 2000 X 30 network. Uneven firing average was observed in the right-most columns of neurons. This simulation had a moderate non-linear to linear conductance ratio (0.38). The histograms contributed by the firing rate of neurons in right-most 10 columns after 8,000 or 10,000 computing iterations (right). **(C)** Without a non-linear component, a neural activity hierarchy doesn’t appear. Absence of non-linear conductance ratio does not naturally lead to the appearance of neuronal priming (left) or a right-skewed activity histogram (right). **(D)** Removing the non-linear component (at 10,000 computational iteration) quickly disrupts an established activity hierarchy and reinstating it (at 20,000 computational iteration) randomly primes a new cohort of neurons. The newly formed hierarchy is clearly distinct from the initial one. **(E-F)** same as **(B-C)**, except for reduced (2%) initial activity levels for **(F)**. Reduced initial neuronal activity (from 6% to 2%) does not lead to neuronal priming. Note the scale of X-axis is different in **(F)**. **(G)** c-Fos expression in the hippocampal CA1 region. (Top) Naïve, mice were not subject to tone or foot shock stimulation; (Bottom) Trained, c-Fos expression 45 min after mice were subject to 7 cycles of trace fear conditioning. **(H)** Histogram of c-Fos expression (normalized to the c-Fos intensity average) in the CA1 45 min after trace fear conditioning. It shows a right-skewed log-distribution and is well fitted with a log-logistic function.

Due to much fewer primed neurons, Ift88 KOs have a much sparser distribution of primed neurons on imaging fields (Fig. 5D). Collectively, these data support the accuracy of primed neuron identification.

### Accumulation of non-linear weighted synaptic transmission naturally leads to activity hierarchy in a simulated neural network

To understand how a small fraction of neurons become more active than others (we term it neuronal priming), we developed a simplified computational model consisting of a two-dimensional array of neurons, in which electrical signals are assumed to propagate from left to right through the array. Each neuron, except in the rightmost column, is assumed to have axon terminals connected to the dendrites of neurons on its right with a variable connection strength *Ski* (see Methods & Fig. 6A). Postsynaptic conductance in the brain is mostly mediated by the AMPA receptors and NMDA receptors ^60^, which we simulated as linear and non-linear weighted synaptic conductances, respectively. Our model also includes a negative conductance, mimicking the inhibition of GABAAR conductance. The simulation was initialized with each neuron being equivalent, other than starting at random initial activation levels. A subset of the left-most neurons was randomly chosen to fire each computational iteration. Neural activity hierarchy naturally developed from left to right, and the right 10 columns were collected to form a firing rate distribution histogram, whereas the left edge was enforced to be random by the simulation. As signal accumulates and the membrane potential depolarizes, neurons in subsequent columns reached the threshold for firing and the iterations were continued to see if neurons with high firing rates would appear. We set the values of the parameters of our model based through experimentation and found that a moderate ratio of non-linear to linear conductance (∼0.38) does produce significant neuronal priming patterns and the resulting firing rates show a right-skewed log-distribution. Interestingly, this ratio is well-aligned with the non-linear to linear conductance ratio in neurons measured by previous studies ^61,62^. Fig. 6B left panel presents the firing rate of the entire neuron array during computational iterations, showing that as the impulse signal progressed from left to right, that preferential pathways appear. The right panel shows the histograms of the firing activity level for the last ten layers on the right edge during iterations, demonstrating a significantly right-skewed log-distribution which were in line with the neuronal activity distribution measured by *in vivo* calcium imaging experiments (Fig. 6B). The non-linear synaptic conductance is required, because exclusion of the non-linear component in the simulation does not lead to the development of neuronal priming over 10,000 computation iterations (Fig. 6C). In addition, removal of the non-linear component eliminates an established neural activity pattern (Fig. 6D), reintroducing it back to the network stochastically causes a new pool of neurons to become more active than others (Fig. 6D).

If the development of neuronal priming has something to do with the non-linear transmission, modeling the NMDAR gate’s opening, then those gates require certain activity levels to open, thus, the low firing activity level will impede neuronal priming and pattern formation. When the patterns do form, the strength of the patterns and the speed at which they form is tied to the firing rate. Indeed, we found that basal neuronal activity levels also affect the progression of neuronal priming. We changed the initial basal activity levels in our simulation from 6% neurons firing in the first layer per cycle to 2% neurons (Fig. 6E-F), and then neuronal priming is not observed, and the distribution of the firing activity is not significantly skewed nor is there much variation in the firing activity levels (Fig. 6F).

To connect our simulation results to the realism of hippocampal memory-eligible neurons, we measured the expression level of c-Fos that reflects immediate early gene expression in response to trace fear conditioning. Trace fear conditioning significantly increased the c-Fos positive neurons in the CA1 region (Fig. 6G). Interestingly, we observed that the expression level of c-Fos displays a similar situation that a small portion of cells with very high signal, whereas other neurons had low expression level. We collected more than 1,600 c-Fos positive CA1 neurons out of 5 control mice to construct a histogram. Similar to the right-skewed distribution of calcium dynamics that measured by SD-of-SD, the c-Fos expression levels demonstrated a right-skewed log distribution that can be fitted with a log-logistic probability density function (Fig. 6H).

### NMDAR blockade by ketamine during trace fear conditioning suppresses general neuronal activity, impairs burst synchronization and associative learning

To experimentally test if NMDAR control neural activity hierarchy in the hippocampus, we combined a pharmacological approach with behavioral test and *in vivo* calcium imaging. We injected mice with 10 mg/kg (a subanesthetic dose) ketamine, an uncompetitive NMDAR antagonist ^39,40^, 5 min before training and subjected them to trace fear conditioning and simultaneously monitored calcium dynamic of hippocampal CA1 neurons. We observed when animals were under the influence of ketamine, they exhibited psychotic behaviors, uncontrolled movement, and radical running (not shown). Repetitive tone foot-shock paired training failed to train animal to associate tone to foot shock (Fig. 7A-B) and animals did not freeze throughout 7 training cycles (not shown). Paradoxically, although animals treated with ketamine moved more radically than animals treated with vehicle, their overall hippocampal neuronal activity was markedly reduced, as shown by the leftward shift of neuronal activity histogram plots and shortened tails in the right (Fig. 7C). Remarkably, when mice were under ketamine, the neural activity hierarchy (or ranking) of hippocampal neurons during conditioning were disrupted (Fig. 7D) and animals could not stably maintain a pool of primed neurons (Fig. 7E). Consistently, ketamine treatment drastically decreased the expression abundance of c-Fos induced by trace fear conditioning (Fig. 7F-G). This data indicates that NMDAR blockade suppresses general neuronal activity, impairs neural synchronization and associative learning.

**Fig. 7.**
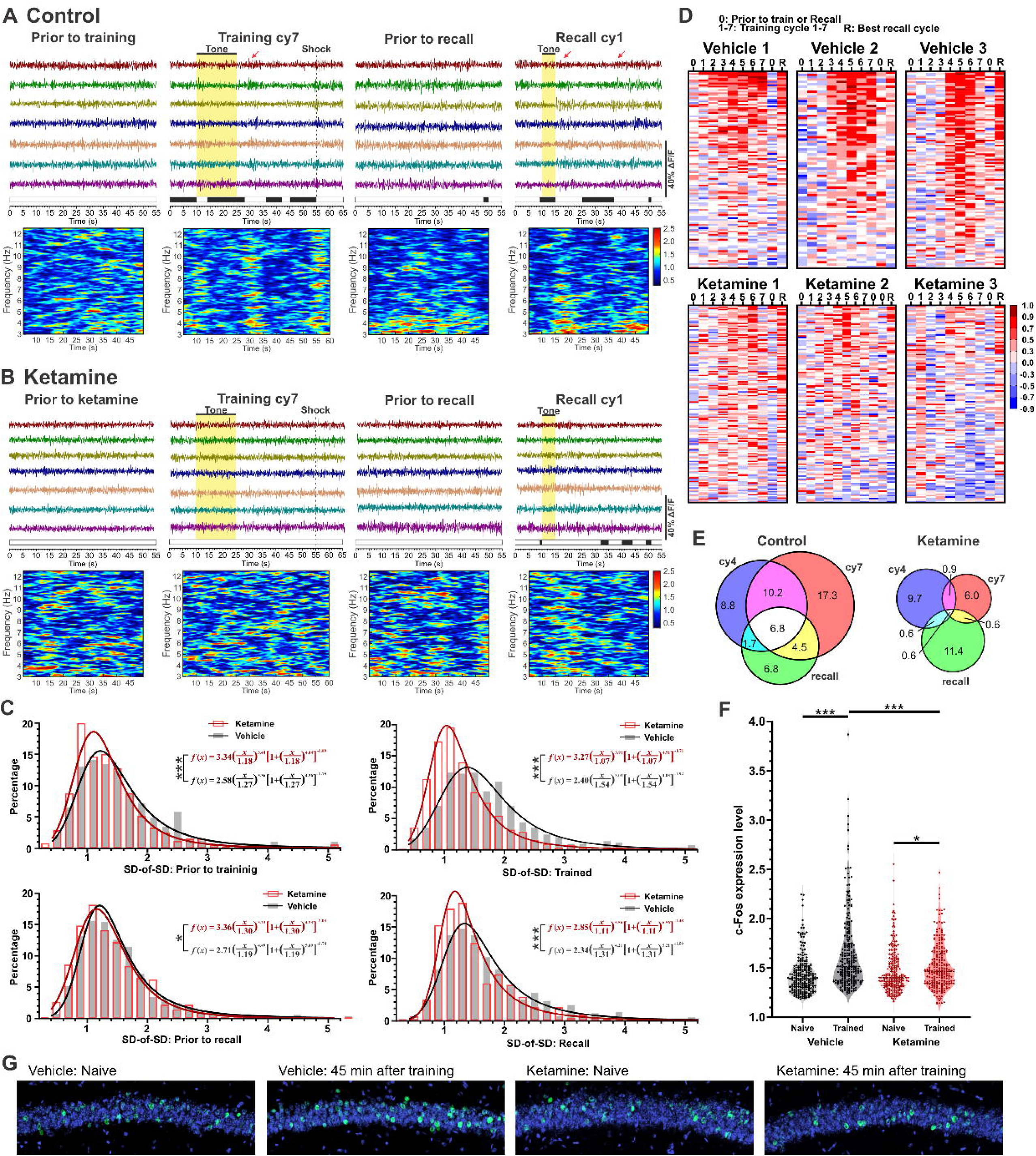
NMDAR blockade by ketamine during conditioning suppresses general neuronal activity and neural synchronization and impairs associative learning. Ketamine (10 mg/kg) or vehicle was injected 10 min before training. **(A-B)** vehicle (A) and ketamine (B) group. (Top) Representative calcium imaging traces of 7 high-activity neurons, filtered by a 1 Hz high-pass Fourier filter. White and black bars denote mouse moving or freezing, respectively. (Bottom) Time-frequency density plots of calcium dynamics of 7 high-activity neurons. **(C)** Activity histograms of all neurons in vehicle control group (grey) and ketamine group (red). The SD-of-SD of individual neurons prior to training and image were acquired at 10 mins after injection (top, left), at the end of training (top, right), prior to recall (bottom, left), and during recall (bottom, right), normalized to the baseline under anesthesia. The fitting curves were made by probability density functions, with K-S tests determining statistical differences. Vehicle group, n = 459 neurons from 4 animals; ketamine group, n = 419 neurons from 5 animals. K-S test between vehicle and ketamine groups, *, p<0.05; ***, p<0.001. **(D)** Pearson correlation coefficient of individual neurons against the major patterns of PCA at different training or recall testing cycles. In the vehicle group (top), the same group of neurons maintain high correlation across various conditions, while in the ketamine group (bottom), a clear hierarchy is not observed. **(E)** Ketamine reduced the percentage of overlapped neurons which were highly correlated with the major pattern at the middle of training (training cycle 4), at the end of training (training cycle 7) and during recall. **(F)** Normalized intensity of c-Fos expression (normalized to the truncated mean) in the CA1 region, 45 minutes after trace fear conditioning. A significant increase in c-Fos expressing was observed in vehicle group after training, which was not observed in ketamine group. Statistical analysis was conducted using unpaired Student t-tests comparing naïve and 45-minutes after training intervals (* p<0.05; *** p<0.001). **(G)** Representative images of hippocampal c-Fos staining under 4 different conditions (with or without ketamine, naïve or 45 minutes after training).

### Loss of neural activity hierarchy contributes to ketamine-induced super-synchronized slow oscillation

Ketamine is known as a dissociative anesthetic ^39,40,42,63^, which is different from isoflurane and other GABAergic anesthetics. We reasoned that through blocking NMDARs, which mediate a higher ionic conductance in primed neurons than silent neurons, ketamine preferentially dampens primed neurons’ activity and thereby flattens neural activity hierarchy (Fig. 8A). We anesthetized mice with ketamine and recorded EEG/EMG. EEG waveforms during awake typically have low amplitude with broad frequency. However, under ketamine-induced anesthesia, EEG waveforms drastically changed to high-amplitude super-synchronized slow oscillation (Fig. 8B). The EEG amplitude under the control of ketamine was much higher than awake (Fig. 8B) and even higher than slow-wave deep-sleep (not shown). While ketamine is a short-acting agent, which has an elimination half-life of ∼13 minutes in mice ^63^, the effect of ketamine on EEG waveforms lasted much longer. Mice did not quickly recover or rapidly restore normal EEG within 100 minutes (Fig. 8B). In contrast, isoflurane initially induced a low-amplitude slow-wave EEG waveform which was mixed with burst suppression, gradually advancing to the isoelectric state (Fig. 8C). After termination of isoflurane exposure, mice recovered quickly, and reinstated normal EEG waveform as awake within a few minutes (Fig. 8C). Clearly, ketamine induces super-synchronized slow oscillation while isoflurane does not. We asked the question if ketamine-induced super-synchronized slow oscillation is partly due to the loss of neural activity hierarchy. To address this, we employed the same neural simulation model (as described in Fig. 6) and included an electric probe in the network to detect neural firing in a local field. We found that the ketamine-induced slow oscillation can be nicely simulated by the model. If the neural network model included the non-linear NMDRA-mediated component, the neural activity detected by the probe was random, low-amplitude and high-frequency (Fig. 8D). However, if removing the non-linear component, the level of general neural activity drastically reduced (Fig. 6D), but neurons in the same layer tends to synchronously fire, which occurred in a very slow phase (Fig. 8F). This simulation mimics the high-amplitude slow oscillation induced by ketamine. Restoring the non-linear NMDAR component de-synchronized neural firing (Fig. 8G). This data suggests that the ketamine-induced super-synchronized slow oscillation is partially caused by the loss of neural activity hierarchy, which neutralizes the differences between primed and silent neurons, allowing them to fire synchronously.

**Fig. 8.**
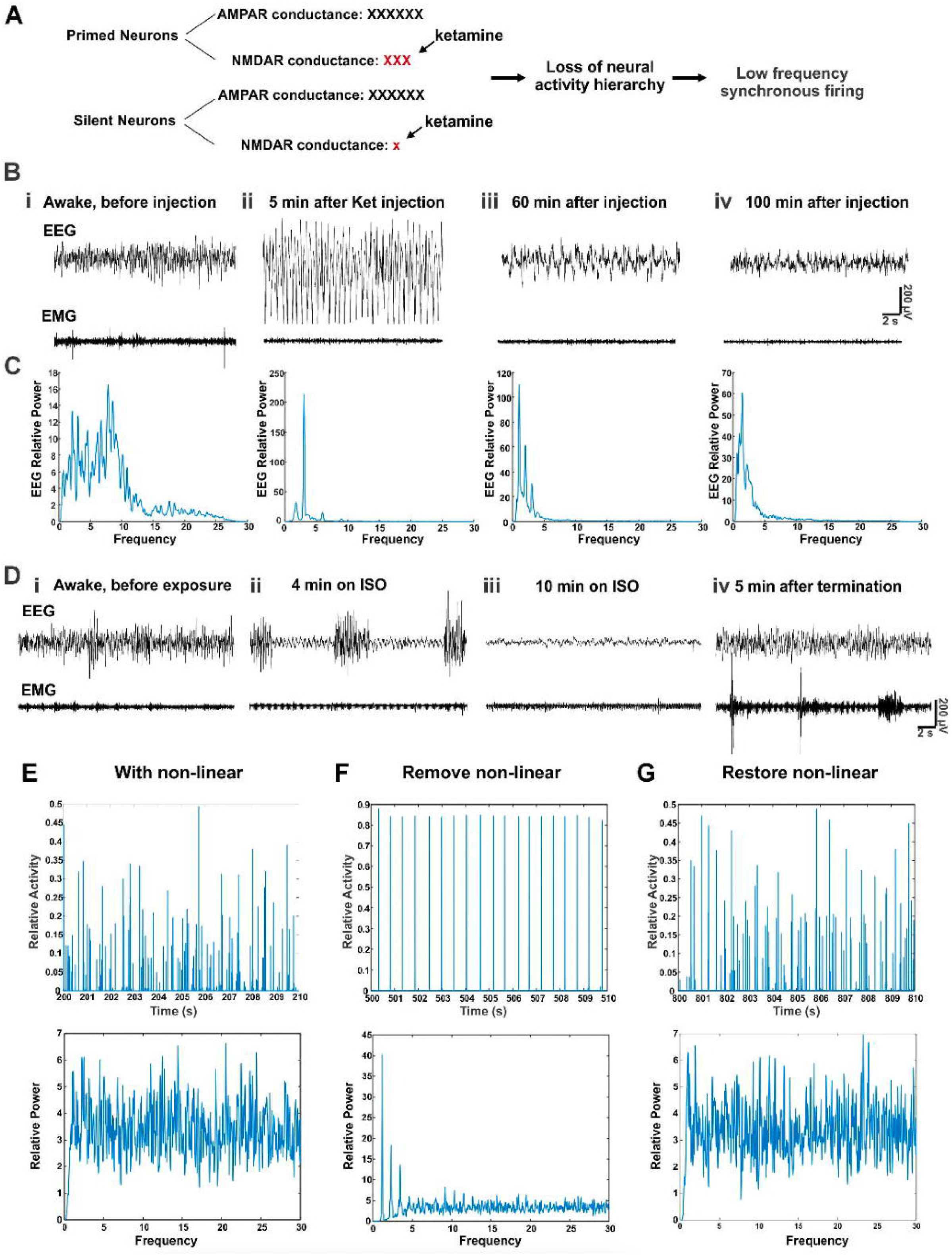
Ketamine-induced super-synchronized slow oscillation can be simulated by our neural network model if the non-linear NMDAR conductance is removed. **(A)** NMDAR-mediated conductance in primed neurons (depolarized neurons) is stronger than that in silent neurons (less depolarized neurons). Ketamine can flatten neural activity hierarchy by predominantly inhibiting NMDAR-conductance of primed neurons. **(B)** Low-frequency, high-amplitude oscillatory waveform induced by ketamine (130 mg/kg) supplemented with xylazine (8.8 mg/kg). (i) EEG during awake (before ketamine injection). (ii-iii) Anesthetic status, 8 min and 18 min after ketamine injection, EEG waveform exhibit low-frequency, high-amplitude oscillation (higher than awake). (iv) EEG waveform did not recover back to normal over 100 min after ketamine injection. **(C)** EEG/EMG waveform induced by 3% isoflurane (ISO). (i) Awake EEG waveform before isoflurane exposure. (ii) Isoflurane-induced EEG burst suppression. (iii) 10 min isoflurane exposure reach the isoelectric state with EEG signals flattened. (iv) A few minutes after the termination of isoflurane exposure, mice quickly recovered to normal and restored normal EEG like awake. **(D-G)** Neural oscillation in simulated network. **(D)** Same simulation as described in Fig. 6, except that an electric probe was included to the network to detect neural activity in a local field. **(E)** The presence of the non-linear component, which ensures the development of neural activity hierarchy, produces randomized neural activity among these neurons. **(F)** Removal of the non-linear NMDAR-conductance, which attenuate overall neuronal activity, generates low-frequency, high-amplitude, super-synchronized oscillation. **(G)** Reinstating the non-linear component leads to randomized neural activity.

## Discussion

This report presents multiple key findings on the activity hierarchy of hippocampal neurons: (1) Based on memory-associated neuronal synchronization and SD-of-SD, we have developed the first numerical method to quantify hippocampal neural activity hierarchy, on top of which we identify primed neurons. (2) A PCA of calcium dynamics has revealed when a trace fear memory is being formed or retrieved, the major pattern of the PCA is predominantly mediated by primed neurons and correlated with mouse freezing behaviors. Conversely, when an animal is not actively engaged in associative learning, the major pattern of the PCA does not fully distinguish itself from other minor patterns. Likewise, cilia KO mice, which exhibit severe learning deficits ^55^, have drastically fewer primed neurons, do not develop trace memory-associated burst synchronization, and repetitive training fails to induce a major pattern that is distinguishable from other minor patterns. (3) In a neural network simulation model that incorporates linear and non-linear weighted components, modeling AMPAR and NMDAR ionic conductance, respectively, primed neurons naturally emerge after iterative computations. The basal activity levels influence the developmental process of neural activity hierarchy. (4) Experimentally, NMDAR blockade by ketamine not only temporally decreases general neuronal activity and impairs associative learning, but also disrupts the existing neural hierarchy. (5) Ketamine-induced super-synchronized slow oscillation can be mimicked through simulation when the non-linear NMDAR component is removed. Together, this work demonstrates that NMDARs control the development and maintenance of neural activity hierarchy, and disruption of the existing hierarchy by NMDAR blockade leads to dissociative and psychosis-like behaviors.

To understand the mechanisms of associative memory, it is necessary to identify memory-eligible primed neurons and track their real-time dynamics when a memory is acquired or retrieved under the physiological condition. However, to date there is a lack of suited numerical method to quantify neural activity hierarchy and identify primed neurons that are actively engaged in associative learning. Approaches relying on c-Fos expression has limitations. For instance, c-Fos can be pathologically activated by unspecific stimuli including neuronal injury ^64^. Moreover, c-Fos is expressed 30-60 minutes after neuronal activation ^3,65,66^, reflecting immediate early gene expression for memory consolidation at the cellular-level. Fos expression does not time-lock with neural dynamics for associative learning or memory acquisition. Additionally, while Fos-based engram cell labeling methods are widely used in manipulating contextual fear memory ^5,67,68^, Fos expression dynamics are not well characterized in trace fear conditioning that requires repetitive training.

Our deep-brain *in vivo* fiber-optic confocal imaging in conjunction with freely behaving trace fear conditioning avoids GRIN lens implantation and pathological neuronal injury, which helps maintain hippocampal neuronal activity very close to the physiological state (Fig. 1). The lateral resolution of the system reaches 3 µm, allowing for neuronal imaging at the single-cell level. We developed multiple empirical methods to ensure that ROI imaging data were collected from single cells (Fig. 1). Therefore, imaging data acquired from this imaging approach is suited for quantifying real-time neural activity hierarchy. After eliminating background patterns or imaging noise, we performed a PCA of all imaging data to extract a major activity pattern that is associated with fear conditioning and then measure the correlation level of individual neurons with the major pattern. The correlation coefficient of individual neurons with the major pattern yields a ranking of hierarchy (Fig. 4). We complemented that method with SD-of-SD (Fig. 5), which clearly differentiates bursting activity from low activity and consistent high-amplitude fluctuation, which also helps distinguish primed neurons from non-primed neurons. The concept of primed neurons, if solely defined by high activity, could be too general, because some high-activity neurons may not be directly engaged in one specific type of learning. Including a factor of memory-associated burst synchronization refines the pool of primed neurons that actively engage in associative learning. We found that SD-of-SD computation aligns nicely with the correlation levels of primed neurons with the major pattern of PCA, because primed neurons are largely clustered in the upper-right corner in the plot of SD-of-SD vs synchronization levels (Fig. 5). This upper-right cluster becomes pronounced with repetitive training, disappears during resting period, and re-appears during successful recall testing. The high correlation of mouse freezing behaviors with the major pattern of primed neurons also support the accuracy of our quantification method (Fig. 3). Further, we marked the location of identified primed neurons on imaging fields (Fig. 5C), showing that primed neurons are not clustered together *in situ*, rather sparsely distributed, and their averaged distance was estimated to be ∼ 40 µm, reminiscent of c-Fos positive neuron distribution in the hippocampus.

This study has yielded an estimation of primed neuron percentage in the mouse hippocampal CA1 region, which approximates to 9.4 ± 3.4% in WT mice. This number is compatible with other estimations ^3,6,69^. However, the percentage of primed neurons in the hippocampus can be affected by numerous factors, including protein expression, gene mutations, animal strains, subject vigilance, and training paradigms etc. If mice are repetitively trained for additional cycles, more neurons will be categorized into primed neurons, as we observed that repetitive training impacts neural dynamic and recruits more neurons to engage in associative learning. This suggests that the pool of primed neurons is not stagnant but subject to dynamical changes. Remarkably, cilia KO mice were found to have lower activity hierarchy. Cilia KO mice have drastically reduced neuronal activity and a much lower percentage (1.9 ± 1.5%) of primed neurons.

Correspondingly, cilia KO mice cannot develop strong burst synchronization (Fig. 2B). This underscores the importance of the burst synchronization of primed neurons to associative learning and memory formation, suggesting that decreased percentage of primed neurons likely accounts for their learning deficits. Cilia KO mice serve as an excellent negative reference to identify primed neurons. The phenotype of cilia KO mice in imaging is in line with our simulation data, showing that reduced basal neuronal activity affects the formation of neural activity hierarchy (Fig. 6). Nevertheless, it is worth mentioning that many proteins may regulate general neuronal excitability, thereby affecting neural activity hierarchy. For example, CREB ^15,45^ and certain type of potassium channels ^22,46,47^ are found to affect neuronal excitability and neuronal allocation for contextual memory formation.

This study underscores an interesting fundamental question: why does the hippocampus need to develop a neural activity hierarchy? There are limited studies that have addressed such a question ^13,16,21,46^. Possible explanations include: (1) A animal subject does not encounter all sorts of sensory stimuli at one time. The nervous system only transduces certain sensory information into the hippocampus for processing. Thus, only a portion of hippocampal neurons become responsive to limited sensory input at specific time. (2) There is a high-density of neurons in hippocampal laminae (DG, CA3 and CA1), which may facilitate associative learning and long-term memory formation. However, it would cause epilepsy if too many neurons in the hippocampus were active simultaneously. Moreover, if all hippocampal neurons had similar levels of high activity, it would cause a huge demand for energy and nutrient support. Glucose or energy supply, glial support such as neurotransmitter synthesis or recycling could not be guaranteed. (3) An animal or human subject constantly acquires new experience or information. Hypothetically it may need a different subset of memory-eligible primed neurons to encode new experiences or memory information. Also hypothetically, there is a rotational mechanism for different pools of hippocampal neurons to take turn to engage in associative memory formation. (4) Neural activity hierarchy is likely required for encoding associative memory information. It enables a selected group of hippocampal neurons to construct an electrical conduit or form unique connectivity ^20^. Additionally, neural activity hierarchy also promotes connectivity preferentially among primed neurons over inactive neurons. Due to elevated excitability, communications among primed neurons are preferentially enhanced so that they could readily develop synchronized activity and synaptic plasticity. (5) Because of synchronized bursting of primed neurons ^20^, encoded information temporally stored in the hippocampus could be more easily entrained to the neocortex for the system-level memory consolidation and long-term storage.

Because neural activity hierarchy facilitates structured information flow (preferentially from primed neurons to primed neurons), maintaining such an activity hierarchy within a certain range must be crucial for sustaining mental health and ensuring intellectual ability. Little is known regarding the developmental and maintaining mechanisms of neural activity hierarchy in the hippocampus and other brain regions. We show for the first time that the NMDAR-mediated non-linear ionic conductance is required for the establishing and maintenance of neural activity hierarchy. In our simulation, the non-linear to linear conductance ratio is a key factor that controls the evolving process of neural activity hierarchy during iterative computation. It is well known that the NMDARs mediate many forms of synaptic plasticity^24,70^. This research has revealed a novel and significant impact of NMDAR on neural activity hierarchy, which is found to emerge naturally through cumulative non-linear NMDAR-mediated synaptic transmission in the network. Importantly, our simulation result on neuronal priming is highly consistent with in vivo imaging and pharmacological study using ketamine. Furthermore, ketamine-induced super-synchronized slow oscillation is likely caused by the loss of neural activity hierarchy, which can be nicely simulated by our neural network model.

One may wonder why ketamine or NMDAR blockade often induces dissociative and psychotropic effects. This study provides critical mechanistic insights into the pathophysiology of psychosis. Because the hippocampus, and other cortical regions as well, has a neural activity hierarchy, a portion of neurons are primed or more active than other neurons. The depolarized primed neurons have a higher portion of postsynaptic NMDAR conductance than those hyperpolarized or non-primed neurons, whose NMDARs are still blocked by magnesium in a voltage-dependent fashion ^48–50^. Ketamine thereby can preferentially block the NMDAR conductance of primed neurons, but not strongly on the NMDARs of non-primed neurons. This biased blockade selectively dampens the activity of primed neurons and eliminate the existing activity hierarchy, suppressing neuronal activities to a uniform lower level. Hierarchy disruption consequently alters neural information flow in the brain, contributing to dissociative and psychotic behaviors. This preferential blockade of NMDARs and hierarchy disruption likely make ketamine a dissociative anesthetic. Moreover, our data also suggests that the status of primed neurons is not permanent but subject to changes. If NMDARs function normally, a neural activity hierarchy can be maintained; if NMDARs are blocked or hypofunctional, neural activity hierarchy cannot be maintained, and the rotation of primed neurons become faster than normal, which consequently impacts neural information processing and leads to psychotic behaviors.

Revealing the mechanism of how neural activity hierarchy is established will not only advance our fundamental understanding of associative learning and cognitive dysfunction-related psychiatric disorders but also catalyze the development of mechanism-based therapies for these disorders. Ketamine and other related NMDAR blockers have been explored for therapeutic usage to treat major depressive disorder and PTSD, particularly for patients who do not respond to conventional treatments ^41,71^. The mechanisms of action of ketamine as a fast anti-depressant are not fully understood. Numerous studies have contributed to generating a variety of explanations ^39–42,72–75^. This study presents a novel mechanistic insight into the therapeutic actions of ketamine in combating major depression: by blocking NMDARs and disrupting the existing neural activity hierarchy in the brain, ketamine and other NMDAR blockers may serve as a “circuit switcher” that reshuffles the ensemble of primed neurons, which consequently interferes with neural information flow in the brain. The “switching” of circuitry not only impacts associative learning, causing psychotic effects, but also can modify the existing neural circuits underlying depression. Thus, ketamine may exert fast-acting anti-depressant effects, at least in part, by re-shaping the existing neural activity hierarchy in the brain. Utilizing this new mechanism of action to disrupt neural activity hierarchy may promote the development of new generation of neurotherapeutics for clinical depression and PTSD. We acknowledge several limitations in this study: (i) Our fiber-optic imaging approach has a low calcium imaging sensitivity. It can detect strong bursting activities but cannot resolve individual calcium spikes. The fiber-optic imaging acquisition system converts weak calcium signals into basal fluctuation. High fluctuation or strong bursting indicates high neuronal activity, while low calcium trace denotes low neuronal activity (not necessarily completely silent). Thus, this limitation does not prevent us from determining neural activity hierarchy. (ii) We used mouse trace fear conditioning as a behavioral paradigm, which is a temporal associated memory, to quantify neural activity hierarchy. Other types of memory-related behavioral paradigms should be employed in future studies to evaluate the generality. (iii) Our neural network model is a fast-scale algorithm and does not incorporate the secondary effects of synaptic plasticity. This limitation warrants future research to evolve this model to increase its realism by incorporating equations for different types of receptors and determine how synaptic plasticity impacts the development of neuronal activity hierarchy.

Together, this presents the first method to quantify real-time neural activity hierarchy in the hippocampal CA1 region, which identifies memory-eligible primed neurons. Our neural simulation indicates that the non-linear NMDAR-mediated synaptic transmission directs the development and maintenance of neural activity hierarchy. We also present experimental data supporting the conclusion that NMDA receptors gate neural activity hierarchy in neural network and loss of control of neural activity hierarchy likely accounts for dissociation and psychosis. Therefore, NMDA receptors not only mediate synaptic plasticity ^24–26^, but also control neural activity hierarchy in the brain to direct neural information flow, engaging in associative learning and sustaining mental health in a novel manner.

## Acknowledgements

This study is supported by NIH Grants K01AG054729, P20GM113131, R15MH126317, R15MH125305, COLE Neuroscience Research Awards, and UNH CoRE PRP awards to XC. YZ and J W are supported by Summer TA Research Fellowships (STAF) and a Dissertation Year Fellowship (DYF) from UNH Graduate School. We thank the University Instrumentation Center for A1R HD confocal imaging service.

## Supplemental Figure Legends

**Fig. S1.**
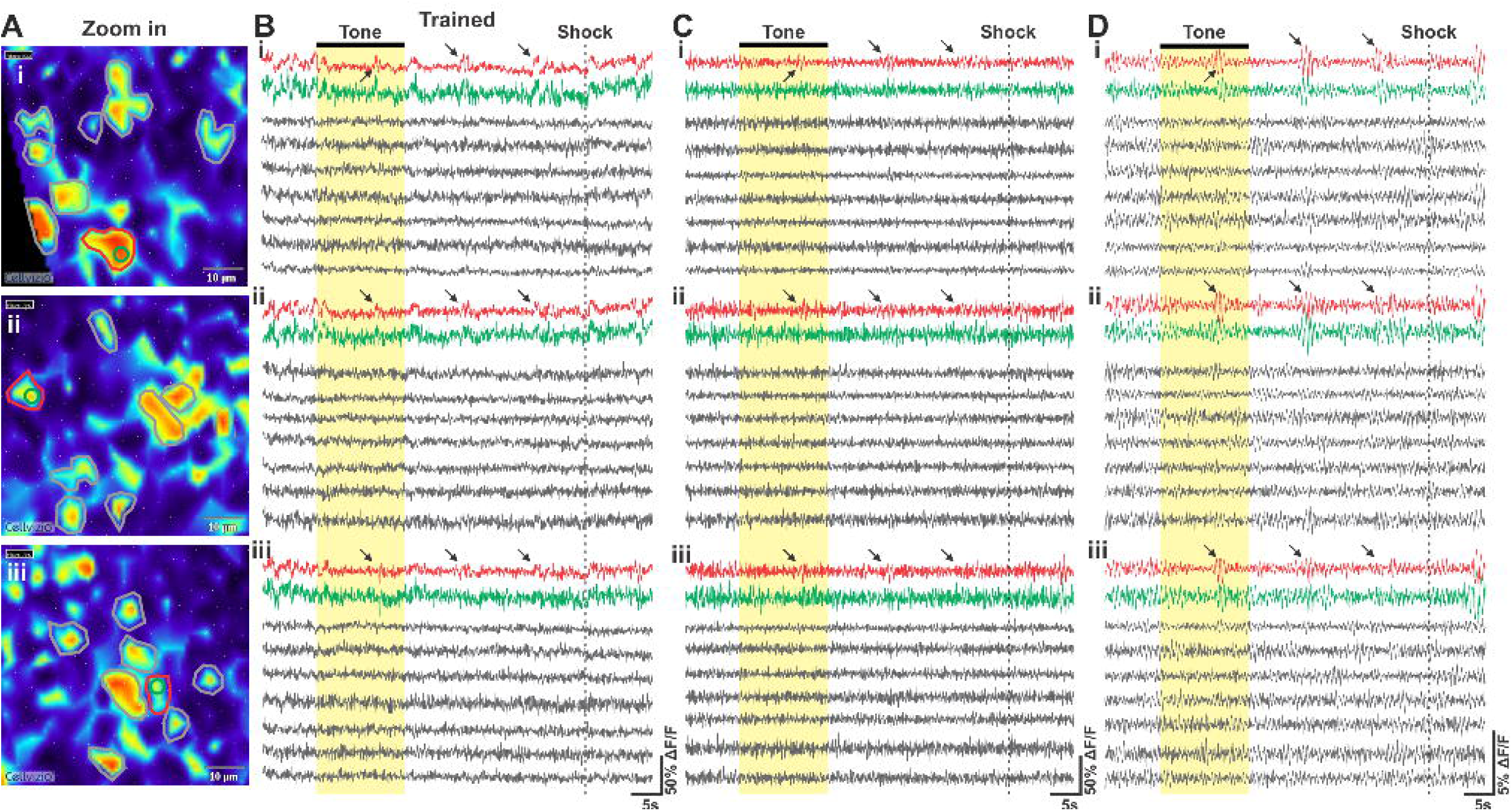
High-pass filter and moving average to show bursts of primed neurons. **(A)** 3 imaging areas, each containing one primed neuron and several non-primed neurons. **(B)** Calcium raw traces of selected neurons (same as Fig. 1I). **(C)** Imaging signals were filtered by a 1 Hz high-pass Fourier filter. **(D)** 1 second moving average of filtered signal. Bursts are highlighted by arrows.

**Fig. S2.**
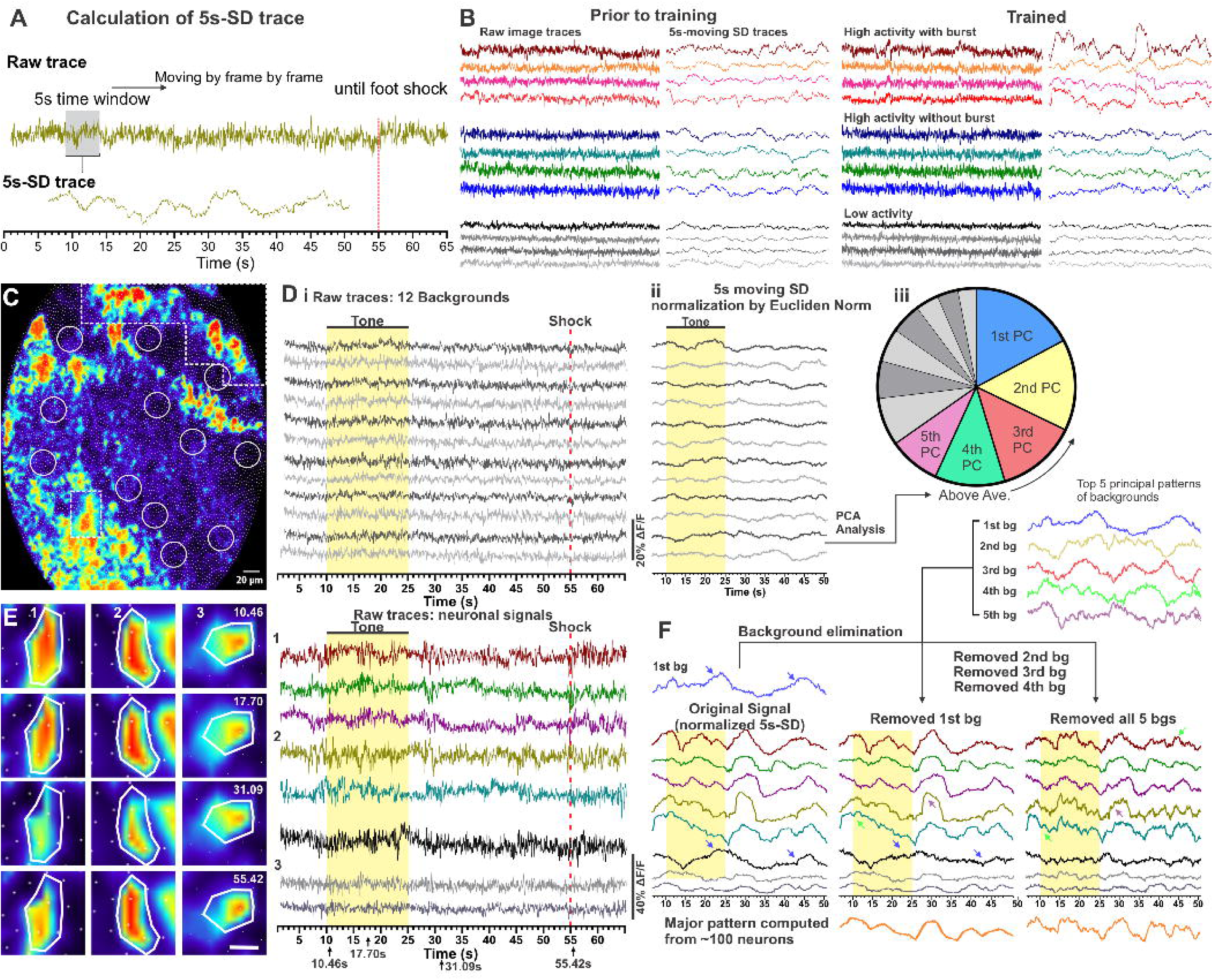
Imaging data pre-processing and background pattern elimination. **(A)** A raw calcium trace was transformed into 5-second moving SD trace. A 5-second window was moving over the image trace frame by frame and computed the SD to generate a 5s-moving SD trace. **(B)** Representative raw calcium image traces (left) and 5s-moving SD traces (right) of high-activity neurons (warm colors), 4 intermediate active neurons (cold colors), and 4 silent neurons (grey). Prior to training (Left) and trained (Right). SD-of-SD of calcium dynamics helps distinguish high-activity neurons from other neurons. **(C)** The Cellvizio confocal fluorescence endoscopic system contains ∼7600 optic-fibers to detect GCaMP6m-emitted fluorescence signals of neurons. Imaging in dashed boxes having different color-contrast setting. 12 background ROIs (grey) were defined as circles covering 50 fibers with minimum GCaMP6 signals to collect background imaging information, which were served as references to eliminate potential motion artifact or background noise. PCA of 12 background ROIs generated principal components for background subtraction. The SD traces were each normalized prior to the PCA. Principal components with PCA levels above the average were used for background noise elimination. **(D)** Representative fluorescence imaging traces of 5 primed neurons (color) and 3 non-primed (grey) neurons were selected. **(E)** Representative time-stamped images of 2 primed neurons (#1&2) and 1 non-primed neurons (#3). Black arrows at the bottom denote the 4 timepoints when images were acquired. **(F)** Neuronal images were first transformed into 5s-moving SD traces. The major background patterns were then used to subtract motion artifacts or background noise step by step. The arrows mark artifacts which were identified and removed during the process of background elimination. Afterwards, a PCA analysis was performed based on signals after background removal. The major pattern obtained from the subsequent PCA is shown on the bottom. The correlation levels of the SD traces of all individual neurons to the major pattern were computed to obtain the synchronization level to identify primed neurons.

**Fig. S3.**
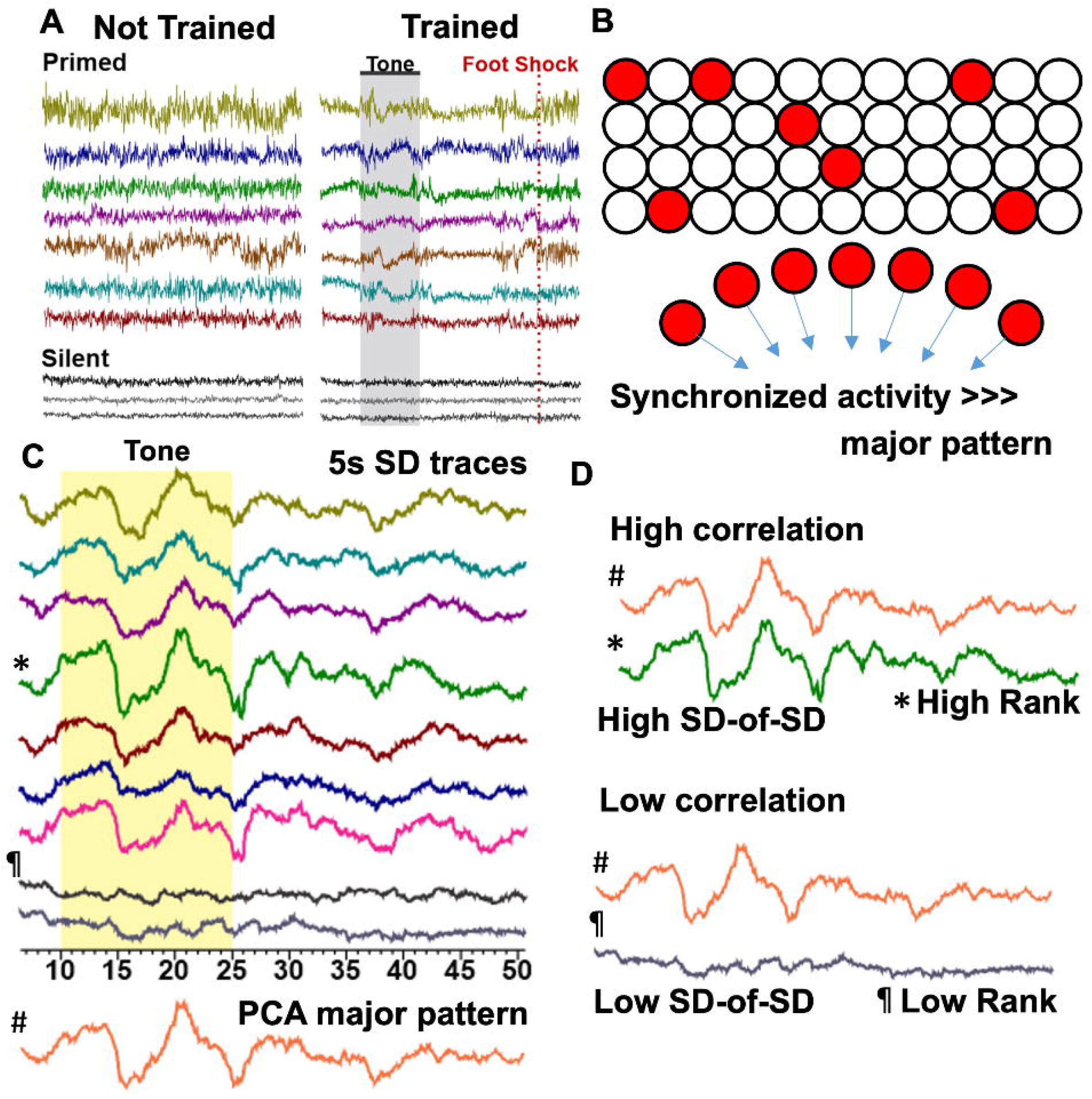
Quantify neural activity hierarchy to identify high hierarchy ranking primed neurons. **(A)** Primed neurons, rather than silent neurons (only showing three), develop burst synchronization during trace fear conditioning. The calcium traces were adopted from our published open-access paper ^20^. **(B)** Primed neurons (red) and silent neurons (white) in the CA1. The PCA major pattern is largely determined by primed neurons’ synchronization. **(C)** 5-second time-windowed standard deviation (5s SD) traces of individual neurons. Color (primed neurons); gray (two silent neurons). The 5s SD traces of all neurons were used in PCA analysis to extract a major pattern (#). **(D)** Individual 5s-SD traces were then used to compare with the major pattern in correlation analysis. The SD of 5s SD traces were also computed to get “SD-of-SD”, which reflects bursting of neuronal activity.

**Fig. S4.**
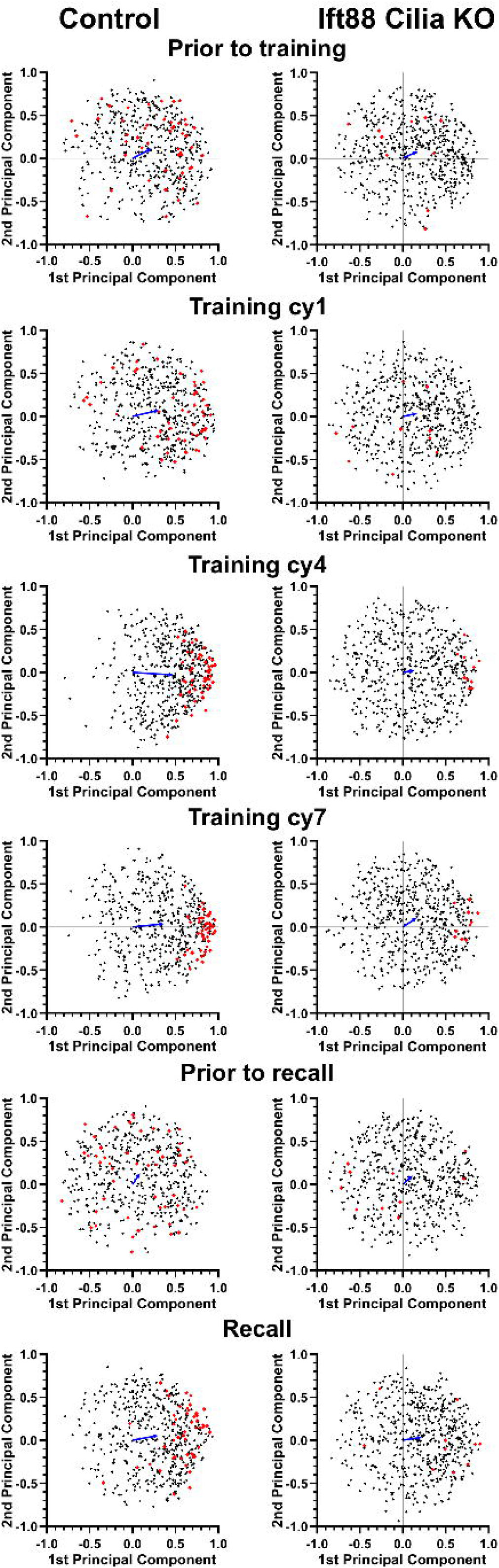
The major pattern of PCA of calcium dynamics is mediated primarily by primed neurons when animals are actively engaged in trace fear conditioning or recall testing. Neuronal contributions to the PCA major pattern during training and recall testing. Controls (Left) and cilia KO mice (Right). Data points are individual neurons’ correlation levels with the principal components (1^st^ and 2^nd^). Data points of X-axis are Pearson correlation coefficient of individual neurons with the 1^st^ principal component. Data points of Y-axis are Pearson correlation coefficient of individual neurons with the 2^nd^ principal component after extracting the 1^st^ component. Arrows pointing from zero to the mean center show the vector addition of two components. In control animals (Left), repetitive training and recall testing increased the weight of the 1^st^ principal component. Individual data points moved to the right, favoring the 1^st^ principal component, while data points of the 2^nd^ principal component remained equally distributed. This change was largely led by primed neurons (labeled in red). In cilia KO animals (Right), individual data points were equally distributed on 1^st^ and 2^nd^ principal components, and training and recall testing did not cause significant changes.

